# Neural plate progenitors give rise to both anterior and posterior pituitary cells

**DOI:** 10.1101/2022.06.12.495791

**Authors:** Qiyu Chen, Dena Leshkowitz, Hanjie Li, Andreas van Impel, Stefan Schulte-Merker, Ido Amit, Karine Rizzoti, Gil Levkowitz

## Abstract

The pituitary is the master neuroendocrine gland, which regulates body homeostasis. It consists of the anterior pituitary/adenohypophysis (AH), which harbors hormones producing cells and the posterior pituitary/neurohypophysis (NH), which relays the direct passage of hormones from the brain to the periphery. It is widely accepted that the AH originates from the oral ectoderm (Rathke’s pouch) whereas the neural ectoderm contributes to the NH. Using single cell transcriptomics of the zebrafish pituitary we characterized *cyp26b1*-positive pituicyte of the NH and *prop1*-positive adenohypophyseal progenitors. We found that these cell types expressed common markers implying lineage relatedness. Genetic tracing revealed that in contrast to the prevailing dogma, neural plate precursors of both zebrafish (*her4.3*+) and mouse (*Sox1+*) contribute to both the neurohypophyseal and adenohypophyseal cells. We further show that pituicytes and *prop1+* progenitors reside in close anatomical proximity and pituicyte-derived RA-degrading enzyme Cyp26b1 fine-tunes differentiation of *prop1+* progenitors into hormone-producing cells. These results challenge the notion that AH cells are exclusively derived from non-neural ectoderm and demonstrate that a cross-talk between neuro- and adeno-hypophyseal cells fine-tunes the development of pituitary neuroendocrine cells.

## Introduction

The pituitary is the master neuroendocrine gland, through which the brain regulates critical physiological functions including metabolism, reproduction, body temperature, water/salt balance, and stress by releasing neurohormones to the circulation. It consists of two major parts: the anterior pituitary also known as the adenohypophysis (AH) and the posterior pituitary also known as the neural lobe or neurohypophysis (NH). The AH mainly contains multiple glandular cells that are distinguished by the hormones they produce (Alatzoglou et al., 2020; Davis et al., 2013; Pearson and Placzek, 2013). The NH is essentially a brain-body interface, which facilitates the passage of the neurohormones oxytocin (OXT) and arginine-vasopressin (AVP) into the periphery blood circulation (Biran et al., 2018; Clasadonte and Prevot, 2018; Murphy et al., 2012). It contains multiple permeable neuro-vascular units formed by hypothalamic axonal projections, fenestrated endothelia and their associated pericytes, and astroglial-like cells, termed pituicytes (Bucy, 1930; Hatton, 1988; Miyata, 2017; Wittkowski, 1986).

The discovery of the dual origin of the pituitary by Rathke in the mid-19^th^ century and its subsequent classification as a gland have been since followed by many studies (Rathke, 1838; Herring, 1908). Thus, the current dogma of pituitary embryonic development manifests that the AH is derived from a non-neural domain, the oral ectoderm (Rathke’s pouch), while the NH is derived from a neural origin as an extension of the third ventricle, the infundibulum, that comes into contact with the adenohypophyseal tissue (Miyata, 2017; Pearson and Placzek, 2013; Rizzoti, 2015).

The doctrine of pituitary development has been further validated by fate mapping experiments in gastrula stage of zebrafish and chick and neurula stage of mouse. These studies have demonstrated that a group of cells within the pre-placodal ectoderm, a horseshoe-like structure that surrounds the neural plate, gives rise to the AH (Couly and Le Douarin, 1988; Dutta et al., 2005; Kouki et al., 2001; Sanchez-Arrones et al., 2017). Although this assertion has been widely accepted, these classical studies relied on labelling techniques that lack single-cell resolution and may have overlooked contributions from other tissues to the adenohypophysis. Indeed, a recent study using transgenic-based lineage tracing has demonstrated that the endoderm also contributes to adenohypophyseal cells in zebrafish (Fabian et al., 2020). Therefore, it is important to re-examine the origins of the anterior and posterior pituitary using more precise genetic and imaging tools. However, despite the physiological importance of the NH, the origin and development of neurohypophyseal cells, and in particular of astroglial-like pituicytes, were impeded by the lack of specific molecular markers and genetic tools.

We have previously employed bulk RNA-Seq of zebrafish pituitary cells, isolated after labelling with a pan-astroglial fluorescent dye, β-Ala-Lys-Ne-AMCA, to characterize the pituicyte-enriched transcriptome, and have validated the expression of top pituicyte markers in the developing pituitary (Anbalagan et al., 2018). Here, using single-cell RNA sequencing, we refined the above-mentioned heterogenous bulk transcriptome thereby revealing the heterogeneity of the zebrafish pituitary. We report the exact molecular signatures of the major cell types, and in particular of the pituicytes and of *prop1-*positive adenohypophyseal progenitors. Using conditional genetic labelling of neural plate cells, we showed that zebrafish *her4.3*-positive neural progenitors, and mouse *Sox1*-positive cells contribute to both adenohypophyseal and neurohypophyseal cells. We further revealed a new cross-talk between pituicytes and *prop1*+ progenitors wherein the pituicyte protein Cyp26b1, which is a retinoic acid (RA) degrading enzyme, attenuates differentiation of *prop1+* progenitors into mature adenohypophyseal cells. Our study challenges the dogma regarding the mutually exclusive embryonic origin of neuro- and the adeno-hypophyseal cells and reveals a new role for pituicytes in refining the differentiation of hormone-producing neuroendocrine cells.

## Results

### Single-cell RNA-sequencing of zebrafish pituitary cells reveal an evolutionarily conserved pituicyte signature

Zebrafish neurohypophyseal tissue is positioned in a pocket-like indentation formed by the adenohypophysis, which makes dissection impossible. We recently exploited the fact that NH cells, including astroglia, actively uptake the fluorescent di-peptide derivative β-Ala-Lys-Nε-AMCA, allowing the isolation and molecular characterization of a pituicyte-enriched cell population (Anbalagan et al., 2018). To better define this population, single cell suspension of β-Ala-Lys-Ne-AMCA-labeled cells from dissected adult pituitaries was sorted into multi-well plates followed by Mars-Seq single-cell sequencing (Jaitin et al., 2014). Pooled data sets from multiple independent samples revealed 1281 pituitary cells that were assigned into nine cell clusters whose annotated cell type identities were determined by comparison to published databases (Figure 1A, B, and Supplementary Table 1). The top differentially expressed marker genes from each cluster are shown in a heatmap and feature plots (Figure 1A, and Supplementary Figure 1A). Notably, the principal pituicyte enriched genes found by scRNA-Seq were consistent with our previously reported bulk RNA-Seq data (Anbalagan et al., 2018). We demonstrated by mRNA *in situ* hybridization that top pituicytes markers obtained from scRNA-Seq of adult tissue (i.e. *cyp26b1*, *cfd,* and *rx3*) were expressed in the developing pituitary at 5-8 days post fertilization (dpf) (Supplementary Figure 1B-D”). In agreement with the assumed astroglial nature of pituicyte (Dellmann and Sikora, 1981; Kawamoto and Kawashima, 1984; Wittkowski, 1986, 1998), known astroglia markers, such as *vim, cx43, cyp19a1b, fabp7a, slc1a3a,* and *slc1a3b*, were enriched in the pituicyte cluster (Figure 1C and Supplementary Table 1).

**Figure 1.**
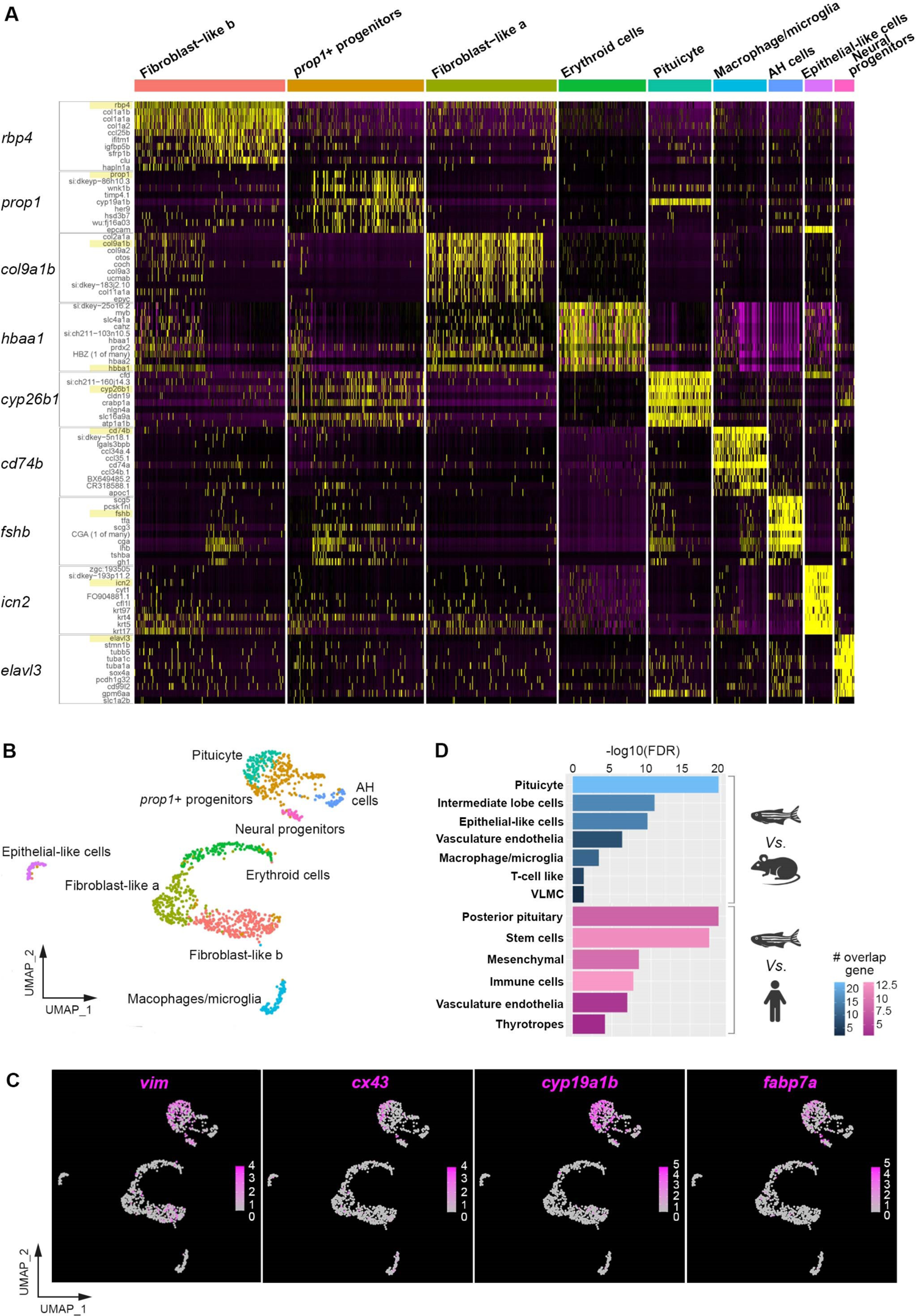
Single-cell RNA sequencing reveals conserved pituicyte molecular signatures with astroglia features. (A) Heatmap showing top ten genes from nine cell clusters of the zebrafish pituitary. Each column represents a single cell with its gene expressions showed in the row. The feature genes for each cluster are highlighted. (B) Uniform manifold approximation and projection (UMAP) depicting various pituitary cell clusters. Designated cell types are color-coded and each dot represents a single cell. (C) UMAP superimposed feature plots illustrating the relative expression of representative pituicyte-enriched astroglial markers (*vim*, *cx43*, *cyp19ab1* and *fabp7a*). The gene expression levels are color-coded with high expression in magenta and low expression in gray. (D) Histogram showing over-representation analyses (ORA) comparing zebrafish pituicyte to scRNA-Seq gene sets of adult mouse neurointermediate pituitary lobe (Chen Q et al., 2020) or human fetus pituitary (Zhang et al., 2020). The bar lengths indicate -log_10_(FDR), and the color-coded scale bars depict the number of overlapping genes.

Many zebrafish pituicyte markers also appeared in our reported single-cell transcriptome of mouse pituicyte (e.g. *Cx43, Dio2, Fabp7, Vegfa, Vim, Rax/rx3* and *Slc1a3*) (Chen Q et al., 2020), as well as in human pituicytes (e.g., *CRABP1, RAX, LHX2, FABP7,* and *SLC1A3*) (Zhang et al., 2020), indicating conservation across vertebrate species. To demonstrate this species conservation in an unbiased manner, we employed over-representation analysis (ORA) (Boyle et al., 2004), which determines whether known biological functions or gene sets are over-represented in an experimentally derived gene list. The ORA comparing the zebrafish scRNA-Seq transcriptome to cell types in the mouse NH (Chen Q et al., 2020) and fetal human pituitary (Zhang et al., 2020) revealed that zebrafish pituicyte gene set was highly enriched in both mouse pituicyte (FDR=1.99E-20) and human fetus posterior pituitary (FDR=6.51E-12) clusters (Figure 1D and Supplementary Table 2). We therefore conclude that the molecular signature of zebrafish, mouse, and human pituicytes is evolutionarily conserved.

### Neurohypophyseal pituicyte and *prop1+* adenohypophyseal progenitors express common markers

Pituicytes and adenohypophyseal cells reside in close anatomical proximity (Anbalagan et al., 2018). Two prominent cell populations we identified in the scRNA-Seq analysis of β-Ala-Lys-Ne-AMCA-labeled cells were *cyp26b1+* pituicyte and adenohypophyseal *prop1+* progenitors (Figure 1 and Supplementary Table 1). *prop1* encodes a paired-like homeodomain transcription factor, which is expressed in progenitors that give rise to AH cells in mammals (Agarwal et al., 2000; Ward et al., 2005). In zebrafish, *prop1* is expressed in the presumptive pituitary placode around 21hpf (Angotzi et al., 2011). We reanalyzed data from a recent, more substantial scRNA-Seq effort from adult pituitary (Fabian et al., 2020) and found that the pituicyte cluster shared 44% of their marker genes (118/268) with *prop1*+ progenitors (Figure 2A). In particular, known radial glia markers such as *cyp19a1b*, *vim*, *slc16a9a*, *slc1a3a*, *fabp7a* and *slit3* (Cosacak et al., 2019; Marin et al., 1989; Menuet et al., 2005; Zhang et al., 2020) were highly enriched in both the pituicytes and *prop1+* cells. We thus envisioned that such markers could serve as a useful tool to study the development of both lineages. In particular, the ventral telencephalic radial glia progenitors marker *slc16a9a* (Cosacak et al., 2019) was highly enriched in both cell types.

**Figure 2.**
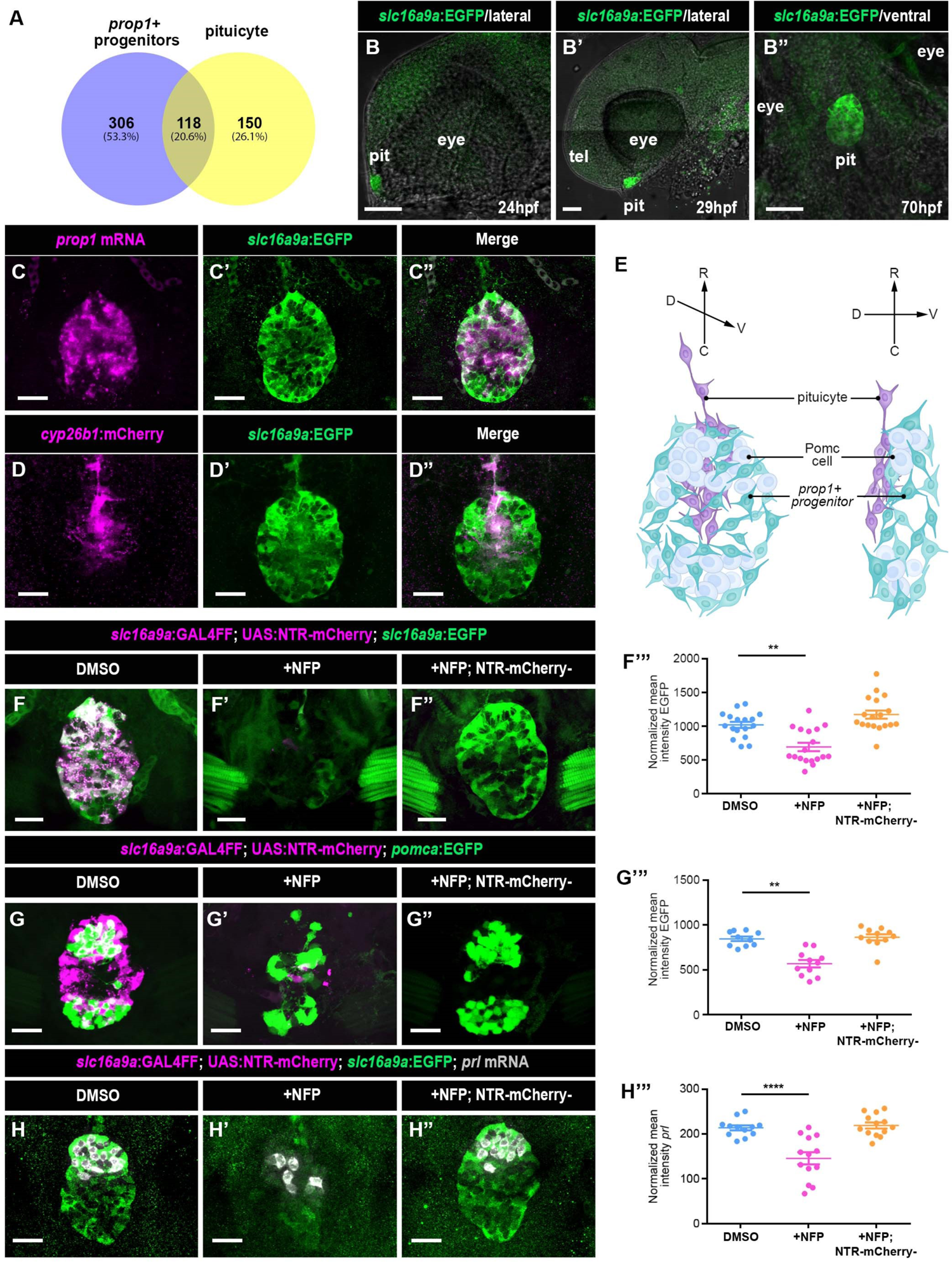
slc16a9a is expressed in the spatially distinct cyp26b1+ pituicyte and prop1+ progenitors. (A) Venn diagram showing the number and percentage of common and differentially expressed genes between *prop1*+ progenitors and pituicyte using re-analyzed whole pituitary scRNA-Seq data by Fabian and colleagues (Fabian et al., 2020). The differentially expressed genes were filtered with average fold change >=2.0 and adjusted *p*-value <=0.05. (B) Developmental expression of the ventral forebrain radial glia reporter Tg(*slc16a9a*:EGFP) in the developing pituitary (**B-B’**) lateral or (**B”**) ventral view, respectively. pit: pituitary, tel: telencephalon. Scale bars, 50μm. (**C-D**) Maximal projected Z-stack of confocal images of transgenic Tg(*slc16a9a*:EGFP) larvae (**C’**, **C”**) following *in situ* hybridization using mRNA probes directed to *prop1* (**C**), or a double transgenic Tg(*cyp26b1*:mCherry*;slc16a9a*:EGFP) (**D’**, **D”**) stained with anti-mCherry antibody (**D, D”**). The *prop1*+ progenitors dispersed in the ventral, while the *cyp26b1*+ pituicyte located in the dorsal central pituitary at 6dpf. Both markers overlap with the Tg(*slc16a9a*:EGFP) signals. Scale bars, 20μm. (E) Schematic illustrations showing the locations of the pituicyte (purple), *prop1*+ progenitors (cyan), and Pomc cells (gray) from a ventral (left) and lateral (right) view of the larval pituitary. C, caudal; D: dorsal; R, rostral; V, ventral. (**F-H**) Representative images and quantifications of genetic NTR-mediated *slc16a9a*+ cell ablation. Triple transgenic Tg(*slc16a9a*:Gal4FF;UAS:NTR-Cherry;*slc16a9a*:EGFP) larvae were used to assess the efficiency of nifuripirinol (NFP)-induced conditional cell ablation (**F**). Larvae were either mock treated with 0.1% DMSO (**F, G**, **H**), or exposed to 5μM NFP which resulted in cytotoxic cell death of nitroreductase (NTR) expressing cells (**F’**, **G’**, **H’**). Sibling larvae with no expression of NTR-Cherry were treated with 5μM NFP to control for non-specific effect of the drug (**F”**, **G”, H”**). Scale bars, 20μm. The scatter plots show the quantifications of normalized intensity of EGFP or mRNA signals in individual larva with the average ± SEM. The respective significance for each treatment is as follows: *slc16a9a*:EGFP (*p*<0.0001 by Kruskal-Wallis test; *p*<0.01 in DMSO vs. NFP, DMSO vs. NFP_Gal4FF/UAS negative is not significant by Dunn’s multiple comparisons test; DMSO n=17, NFP n=18, NTR-mCherry-negative n=18. **F’”**), *pomca*:EGFP (*p*=0.0002 by Kruskal-Wallis test; *p*<0.01 in DMSO vs. NFP, DMSO vs. NFP_Gal4FF/UAS negative is not significant by Dunn’s multiple comparisons test; DMSO n=10, NFP n=11, NTR-mCherry-negative n=11. **G’”**), and *prl* mRNA (*p*<0.0001 by one-way ANOVA test; *p*<0.0001 in DMSO vs. NFP, DMSO vs. NFP_Gal4FF/UAS negative is not significant by Dunnett’s multiple comparisons test; DMSO n=12, NFP n=13, NTR-mCherry-negative n=13. **H’”**). At least three independent replicates were conducted for each group.

We next generated transgenic reporters [Tg(*slc16a9a*:EGFP) & Tg(*cyp26b1*:mCherry)] that were found to faithfully represented the endogenous expression of *slc16a9a* (pituicyte and *prop1*+ progenitors marker) and *cyp26b1* (pituicyte marker) mRNA (Figure 2B and Supplementary Figure 2A-B). *slc16a9a*:EGFP was expressed in the pituitary anlage as early as 24 hours post fertilization (hpf) and maintained its specific expression during development through adulthood with weak expression in the ventral telencephalon and the inner ears (Figure 2B and Supplementary Figure 2C-E). At 5-6dpf, *slc16a9a:*EGFP co-expressed with *prop1* mRNA as well as with Tg(*cyp26b1*:mCherry) reporter. *prop1*+ progenitors were located in the ventral pituitary, whereas the *cyp26b1+* pituicytes resided in the dorso-medial pituitary abutting *prop1+* cells of the tissue (Figure 2C-E and Movie 1). Of note, *cyp26b1* and *prop1* did not overlap (Figure 1A&B, Supplementary Table 1, Figure 2C-E, and Movie 1).

Although *slc16a9a* was highly expressed in *prop1+* progenitors, it was not found in differentiated adenohypophyseal hormones-producing corticotropes (Pomc) cells (Supplementary Figure 2F-G). However, we reasoned that the differentiation of *prop1+* cells towards Pomc cells might involve an intermediate *slc16a9a*+*prop1*+*pomc*+ population. In support of this assumption, we found that ablation of *slc16a9a+* cells affected the development of differentiated AH cells (Figure 2F-H). This was demonstrated using a Tg(*slc16a9a*:Gal4FF;UAS:NTR-mChery) system, in which cytotoxic death of nitroreductase (NTR) expressing cells was induced by nifuripirinol (NFP) (Bergemann et al., 2018; Curado et al., 2008). Thus, embryos were treated with NPF between 3-4 dpf and thereafter allowed to recover for 48 hours to reduce possible acute inflammatory responses caused by the cell death. The average efficiency of *slc16a9a+* ablation was determined to be 32.3% by quantifying the remaining normalized EGFP signals from the surviving *slc16a9a*+ cells in the triple transgenes Tg(*slc16a9a*:Gal4FF; UAS:NTR-mCherry; *slc16a9a*:EGFP) (Figure 2F). Ablating the *slc16a9a*+ cells reduced the mean intensity of Pomc cells and *prl* mRNA signals by 32.8% and 31.9%, respectively with no effect on Gal4FF/UAS-negative siblings (Figure 2G-H).

These results reveal that the pituicytes and *prop1*+ progenitors are closely related cell types, which express common radial glia markers. Furthermore, *slc16a9a* is a novel common marker for *cyp26b1*-positive astroglial pituicyte of the NH and *prop1*-positive progenitors that give rise to hormone-producing adenohypophyseal cells.

### Neural precursors contribute to both neurohypophyseal and adenohypophyseal cells

The developmental origin of pituicytes is still unknown since their first anatomical characterization nearly a century ago (Bucy, 1930). This is mainly due to the lack of precise genetic tools for the study of pituicytes development in both mammals and fish. The close anatomical proximity and shared radial/astro glia markers of pituicytes and *prop1+* adenohypophyseal progenitors implied developmental association between these two cell lineages (Figure 1 and 2, Supplementary Table 1). This prompted us to explore the progenies of neural plate cells at a single cell level using a genetic tracing approach allowing precise spatial-temporal control (Figure 3A).

**Figure 3.**
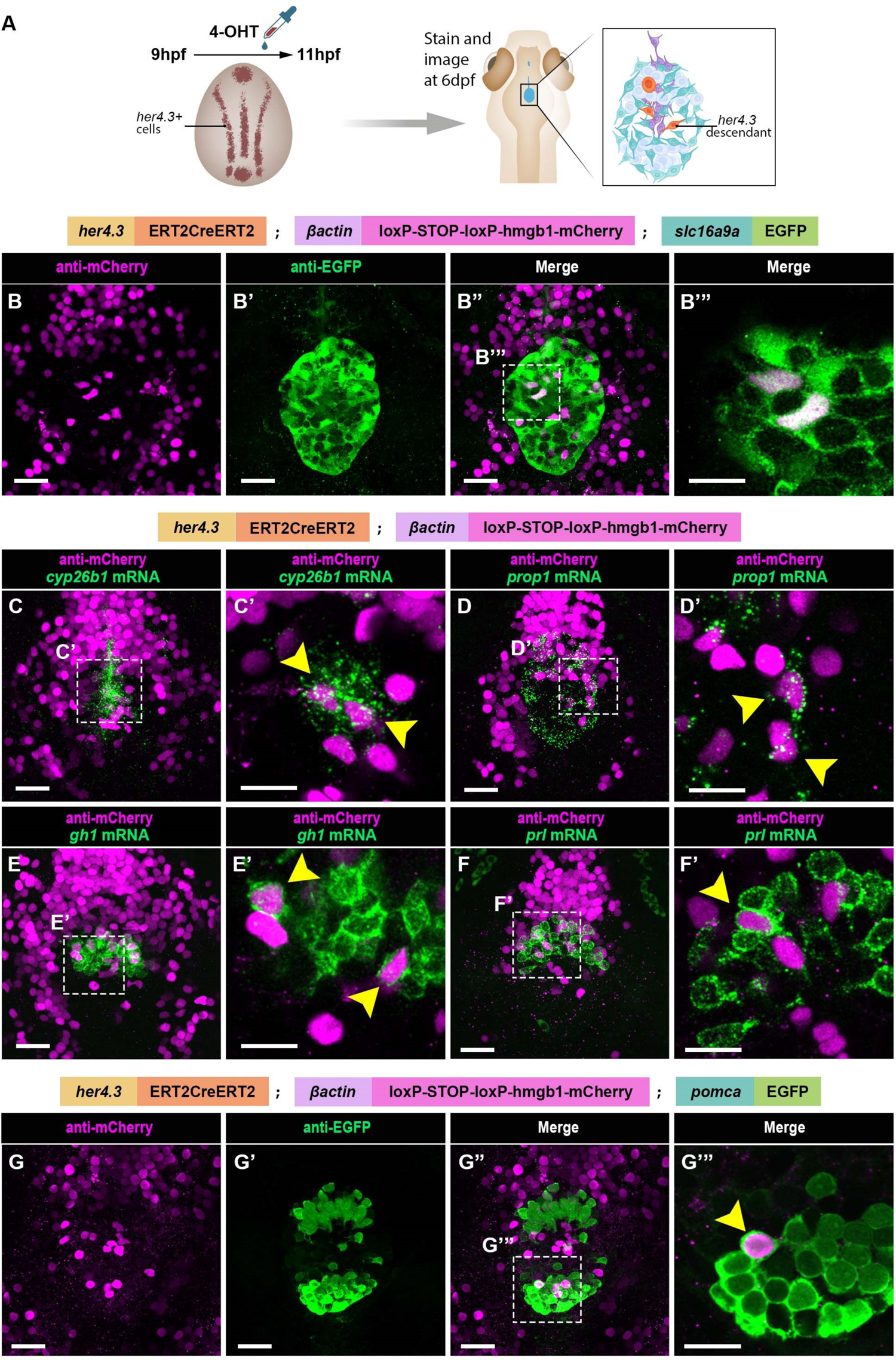
Zebrafish neural plate her4.3-positive progenitors contribute to both NH and AH cells. (A) Schematic representation of the conditional lineage tracing procedure of pituitary cells visualized in **B-G**. Embryos harboring a tamoxifen-inducible transgenic Tg(*her4.3*:ERT2CreERT2;*βactin*:loxP-STOP-loxP-hmgb1-mCherry), which drives nuclear mCherry expression in *her4.3* progenitors, were exposed to 10μM 4-hydroxytamoxifen (4-OHT) at 9 hours post fertilization (hpf) for 2h, followed by extensive washing. Thereafter, embryos were incubated at 28°C until 6dpf for staining and imaging. (B) Confocal Z-stack image of a triple transgenic larva: Tg(*her4.3*:ERT2CreERT2;*βactin*:loxP-STOP-loxP-hmgb1-mCherry;*slc16a9a*:EGFP), wherein neural plate *her4.3* decedents and *slc16a9a*+ cells were labeled as described in **A**, and visualized by co-immunostaining anti-mCherry (**B**) and anti-EGFP (**B’**) antibodies. The merged fluorescent channels of a single confocal Z-plane image (**B’”**) in the selected region (white dash square in **B”**) show overlapping *slc16a9a*:EGFP and *her4.3:*mCherry cells. Scale bars, 20μm in **B-B”** and 10μm in **B’”**. (**C-F**) Transgenic Tg(*her4.3*:ERT2CreERT2;*βactin*:loxP-STOP-loxP-hmgb1-mCherry) larvae were treated as descried in **A,** followed by *in situ* hybridization with HCR-based probes directed to (**C-C’**) *cyp26b1*, (**D-D’**) *prop1*, (**E-E’**) *gh1* and **(F-F’**) *prl*. Scale bars, 20μm. High-magnification single z-plane images of the selected (dashed square) regions are shown in **C’, D’, E’** and **F’** in which cells with overlapping signals were marked by yellow arrow heads. Scale bar, 10μm. (**G**) Neural plate *her4.3* decedents visualized as above in the triple transgenic Tg(*her4.3*:ERT2CreERT2;*βactin*:loxP-STOP-loxP-hmgb1-mCherry;*pomca*:EGFP) showing contribution of *her4.3*+ progenitors to adenohypophyseal *pomc* cells. Panel **G**”’ is a single Z-plane image of the selected (dashed square) region in **G”** demonstrating double-positive *pomca*:EGFP;*her4.3:*mCherry. Scale bars, 20μm in **G-G”,** 10μm in **G’”**.

We labeled *her4.3+* cells in the neural plate using a triple transgenic fish containing *her4.3*:ERT2CreERT2 (Boniface et al., 2009), the ubiquitous CRE-dependent reporter: *Tg(βactin*:LOXP-STOP-LOXP-hmgb1-mCherry) (Wang et al., 2011) and the Tg(*slc16a9a*:EGFP) reporter (Figure 3A,B-B’”). *her4.3* is a Notch target that is expressed by neural precursors in the neural plate (Takke et al., 1999). Embryos were treated with 4-hydroxytamoxifen (4-OHT) for 2 hours, shortly after the neural plate was formed (9 to 11hpf) and much before *prop1, slc16a9a* and *cyp26b1* are expressed in the pituitary primordium [20-24hpf; (Angotzi et al., 2011; Toro et al., 2009)]. No CRE-ERT2 activity was detected in the absence of 4-OHT (Supplementary Figure 3). The progenies of permanently labeled *her4.3+* neural plate cells were thereafter traced in the pituitary at 6dpf (Figure 3A). Using this treatment condition, we found that multiple *scl16a9a*+ cells were derived from the *her4.3* lineage (Figure 3B). As expected, based on their radial glial identity, *cyp26b1*+ pituicyte (7 of 8 larvae) were found to be descendants of *her4.3+* neural plate cells (Figure 3C). Surprisingly, a subset of *prop1*+ AH progenitors (8 of 9 larvae) were also labelled, indicating their neuroectodermal origin (Figure 3D). Consistently, neural plate derived *her4.3* cells gave rise to subsets of the differentiated AH cells including those producing growth hormone (*gh1*) (9 of 10 larvae), prolactin (*prl*) (4 of 9 larvae) and proopiomelanocortin (*pomc*) (5 of 12 larvae) (Figure 3E-G). These observations manifest the common neural origin of *slc16a9a;*c*yp26b1*+ pituicyte, and *slc16a9a;prop1+* progenitors of hormone-producing AH cell types.

To demonstrate evolutionary conservation of our findings, we examined the contribution of neural plate progenitors to both anterior and posterior lineages of the mouse pituitary. We performed lineage tracing of *Sox1*-positive progenitors using the *Sox1^CreERT2^;Rosa26^RTomato/+^* mouse (Kicheva et al., 2014). SOX1 is an early developmental factor, which starts to be expressed at the time of neural induction and promotes neural fate acquisition (Pevny et al., 1998). We induced *Sox1*^CreERT2^ activity by means of a single injection of tamoxifen at 6.5dpc and harvested embryos later at either Rathke’s pouch stage (11.5dpc) or late gestation (18.5 and 19.5dpc). We found that *Sox1*+ progenitors contribute to both Rathke’s pouch (4 of 9 embryos) and the ventral diencephalon (8 of 9 embryos) (Figure 4A). Embryos harvested at late gestation displayed contribution of 6.5dpc-induced *Sox1CreERT2;Rosa^26RTomato/+^*neural progenitors to almost all endocrine lineages of the anterior lobe including, LH, GH, TSH and POMC (Fig. 4B-E) as well as *Sox2*+ stem cells (Figure 4F). Labelled cells were also observed in the posterior lobe of the pituitary and, as expected, some SOX1 descendants represented pituicytes as shown by the expression of LHX2 (Figure 4G)(Chen Q et al., 2020; Zhao et al., 2010). These results show that *Sox1* positive neural plate progenitors contribute to both anterior and posterior pituitary of the mouse. Taken together, the analyses of neural plate lineages in zebrafish and mouse challenge the dogma regarding the strictly separate embryonic origin of the NH and the AH.

**Figure 4.**
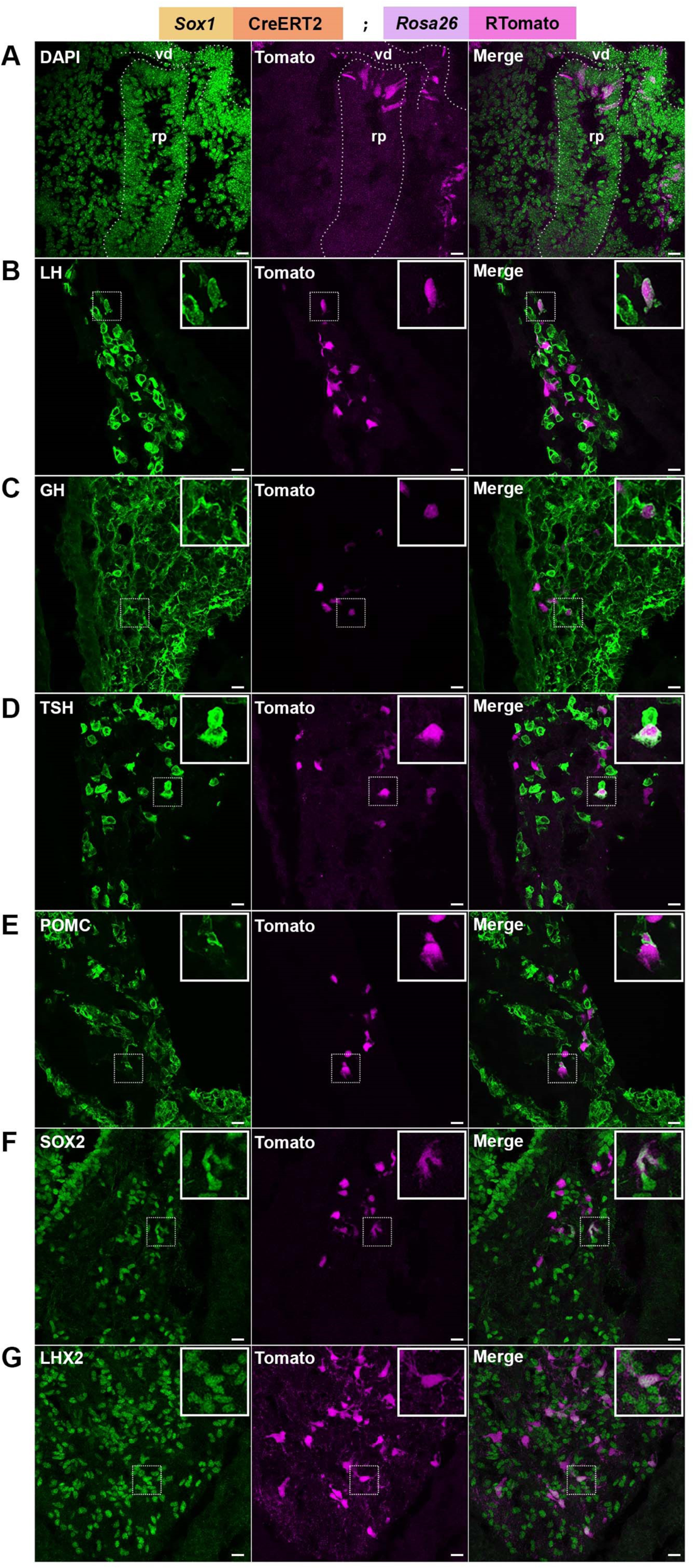
Mouse neural plate *Sox1-*positive progenitors contribute to both NH and AH cells. (**A**) A sagittal section of a 11.5dpc *Sox1^CreERT2/+^*; *Rosa26^RTomato/+^* mouse embryo induced with Tamoxifen at 6.5dpc with DAPI staining and tomato fluorescence. The tomato positive cells are seen both in Rathke’s pouch (rp) and developing ventral diencephalon (vd). Both regions are indicated with dashed lines. Scale bars, 15μm. (**B-G**) Immunofluorescent staining for pituitary markers on sections from 18.5dpc or 19.5dpc *Sox1^CreERT2/+^*; *Rosa26^RTomato/+^* pituitaries from embryos induced at 6.5dpc. Tomato fluorescence was detected in (B) gonadotrophs (LH positive), (C) somatotrophs (GH positive), (D) thyrotrophs (TSH positive), (E) corticotrophs (POMC positive), (F) stem cells (SOX2 positive) and (G) pituicytes (LHX2 positive). Scale bars, 15μm.

### Pituicyte-derived Cyp26b1 attenuates differentiation of *prop1+* progenitors into mature adenohypophyseal cells

The neighboring location of *slc16a9a;cyp26b1*+ and *slc16a9a;prop1*+ cells also implied a functional crosstalk between pituicytes and adenohypophyseal progenitors. It was shown that retinoic acid (RA) promotes the transition of *prop1*+ progenitors into hormone-producing cells. Prop1 is necessary to activate the RA signaling pathway in a cell-autonomous manner through the induction of the RA-synthesizing enzyme Aldh1a2 (Cheung and Camper, 2020; Cheung et al., 2018; Yoshida et al., 2018). Since the pituicyte factor *cyp26b1* encodes a RA degrading enzyme, we hypothesized that the mutually exclusive expression of Cyp26b1 and Prop1 in neighboring pituitary cells fine-tunes adenohypophyseal differentiation by locally restricting RA signaling to *prop1+* progenitors. Thus, we expected that inhibition of Cyp26b1 will promote adenohypophyseal cell differentiation.

To address this, we first tested this hypothesis using *cyp26b1*/*stocksteif* mutants (Spoorendonk et al., 2008) and found a significant increase of the *prop1*+; *pomca*:EGFP+ proportion in the *cyp26b1^(–/–)^* larvae compared to their wild type siblings (Figure 5A, C-E). To further substantiate this result, we blocked Cyp26b1 enzymatic activity using a specific Cyp26 inhibitor, R115866 (a.k.a. talarozole), allowing temporally-controlled inhibition of the enzyme and avoiding possible early developmental effects (Nelson and Isoherranen, 2013). This analysis showed that R115866 treatment between 4 to 5dpf led to a similar increase in *pomca/prop1* colocalization (Figure 5F), as well as of *pomca*:EGFP+; *slc16a9a*:mCherry-NTR+ cells over the total *slc16a9a*:mCherry+ cells (Figure 5B&G). No overlapping signal between *cyp26b1:*mCherry+ pituicyte and *pomca*:EGFP+ cells was found with or without R115866 treatment or in the *cyp26b1^(–/–)^* embryos suggesting a non-cell autonomous effect of pituicyte-derived Cyp26b1 on the differentiation of *prop1+* progenitors to pomc cells (Supplementary Figure 4). Taken together, we provide a new level of understanding into the lineage relationships and functional cellular cross-talk between posterior and anterior pituitary cells.

**Figure 5.**
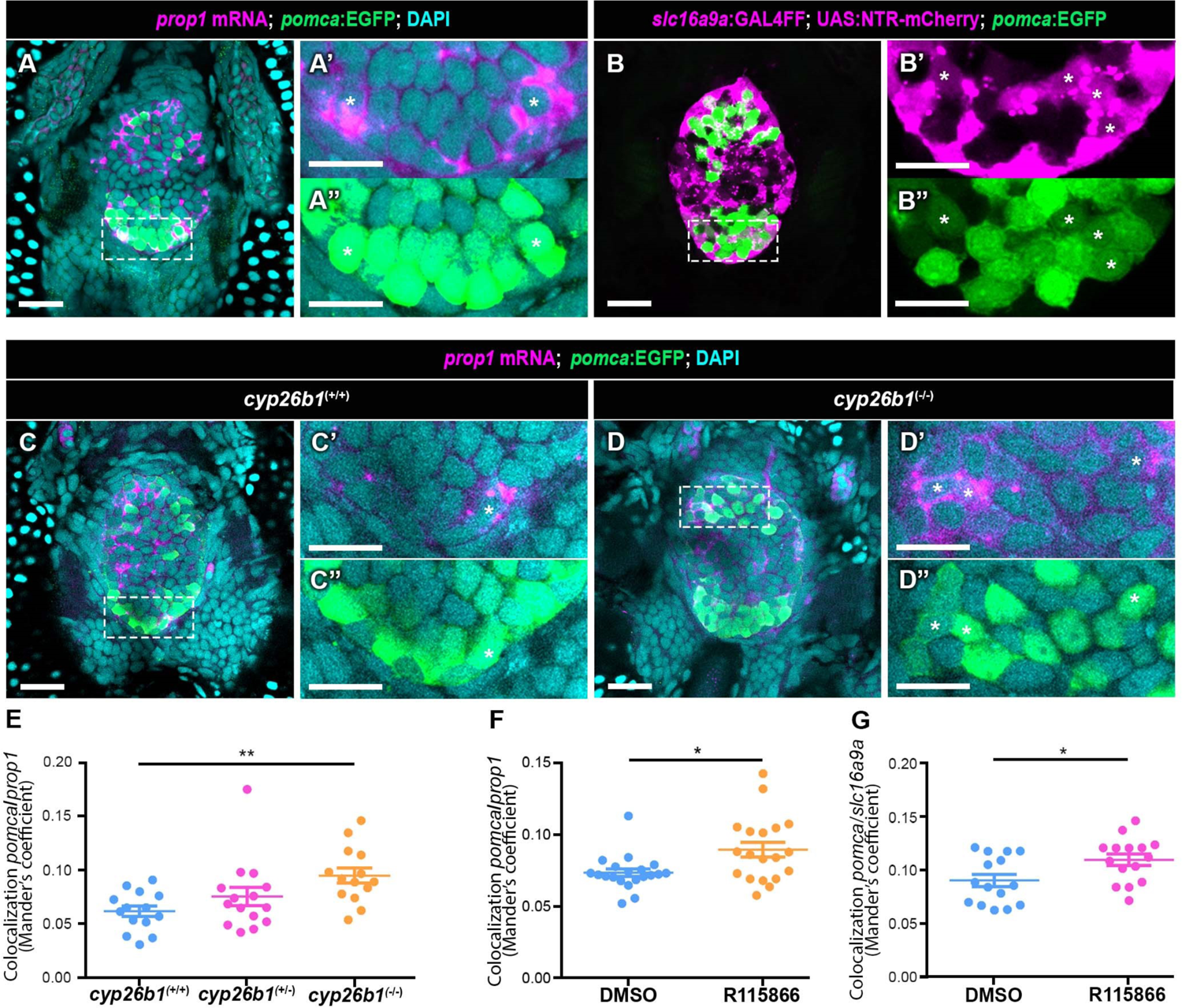
Loss of function of cyp26b1 drives differentiation of prop1+ progenitors. (**A-D**) Single confocal Z-plane images of the pituitary in 5dpf harboring either green Tg(*pomca*:EGFP) (**A**, **C**, **D**) or red Tg(*slc16a9a*:GAL4FF;UAS:NTR-mCherry) (**B**) reporters. Larvae were subjected to *prop1* mRNA in situ hybridization followed by anti-EGFP immunostaining (**A**, **C**, **D**) or imaged for mCherry and EGFP fluorescence (**B**). Samples were subjected to DAPI staining to mark nuclei. High-magnification images of the white dashed boxes show overlapping *prop1+*;*pomc*+ cells (**A, C, D**) or *slc16a9a+*;*pomc*+ cells (**B**) that are marked by white asterisks. *cyp26b1^(+/+)^* (**C**) and *cyp26b1^(-/-)^*(**D**) were determined by genotyping before staining. Scale bars 10μm, expect for 20μm in **A,B,C** & **D**. (**E-G**) Quantification of the overlapping Pomc and *prop1* over total *prop1* signals by Mander’s coefficient revealed significant increase in (**E**) *cyp26b1^(^*^-/-)^ compared to *cyp26b1^(^*^+/+)^ larvae and in (**F**) 2μM R115866 treated larvae compared to the DMSO control. (**G**) Mander’s coefficient of Pomc/*slc16a9a* significantly increased in the 2 μM R115866 treated larvae comparing to DMSO controls. Scatter plot showing individual larva with average ± SEM. (**E**) *p*=0.0035 by Kruskal-Wallis test; *p*<0.01 in *cyp26b1*^(+/+)^ vs. *cyp26b1*^(-/-)^, *cyp26b1*^(+/+)^ vs. *cyp26b1*^(+/-)^ was not significant by Dunn’s multiple comparisons test. *cyp26b1*^(+/+)^ n=14, *cyp26b1*^(+/-)^ n=15, *cyp26b1*^(-/-)^ n=14. (**F**) *p*=0.0109 by Unpaired *t*-test with Welch’s correction, DMSO n=20, R115866 n=19. (**G**) *p*=0.0212 by Unpaired *t*-test with Welch’s correction, DMSO n=15, R115866 n=15. At least three independent replicates were conducted for each group.

## Discussion

The pituitary is an evolutionarily conserved neuroendocrine organ of significant physiological importance. It has been well accepted that its main tissue components, the AH and the NH, originate from the embryonic oral ectoderm and the neural ectoderm, respectively (De Beer, 1924; Herring, 1908; Sanchez-Arrones et al., 2017). Here, we revealed that in contrast to the prevailing dogma, neural plate progenitors of both zebrafish and mouse contribute to both adenohypophyseal and neurohypophyseal cell types. We further demonstrate a new cellular cross-talk between neurohypophyseal pituicyte cells and adenohypophyseal progenitors by showing that the pituicyte-specific factor Cyp26b1 constitutes a critical regulator of the transition of adenohypophyseal *prop1*-positive progenitors into hormone-producing cells.

### Single-cell molecular analysis of pituicytes

The intricate cellular heterogenicity of the vertebrate pituitary was recently revealed by us and others using scRNA-Seq analysis of zebrafish, mouse, human and rat pituitary (Chen Q et al., 2020; Cheung et al., 2018; Fabian et al., 2020; Fletcher, 2021; Mayran et al., 2019; Zhang et al., 2020). However, defining the exact molecular signature of zebrafish neurohypophyseal cell types, and in particular the pituicytes, has been lacking. The single cell transcriptome reported herein refines our previous bulk sequencing analysis (Anbalagan et al., 2018) and highlights the exact pituicyte-specific molecular identity. Thus, we identified specific markers for pituicyte, *prop1*+ progenitors, neural progenitors, macrophage/microglia, and fibroblasts which also appeared in the transcriptome of enriched pituicyte population identified by bulk RNA-Seq analysis (Anbalagan et al., 2018).

In agreement with our reported mouse pituicyte transcriptome (Chen Q et al., 2020), zebrafish pituicytes express bonafide astrocyte/radial glia markers (e.g. *slc1a3a, slc1a3b, vim, cx43, cyp19a1b,* and *fabp7a*) as well as tanycyte markers (e.g. Rax/rx3). This further exemplifies the notion that vertebrate pituicytes belong to a common radial glia lineage (Clasadonte and Prevot, 2018; Rodríguez et al., 2019). Notably, although the top differentially-enriched markers defining pituicytes of mouse, human and zebrafish were not identical, the evolutionary conservation between the three species was evident at the transcriptome level. This was demonstrated by an unbiased bioinformatic ORA analysis (Boyle et al., 2004) of the entire transcriptome of zebrafish pituicytes compared to that of mouse (Chen Q et al., 2020) and human (Zhang et al., 2020).

We have recently reported that commonly used marker genes, which have been so far considered as ‘pituicyte-specific’ display promiscuous cell-type specificity (Chen Q et al., 2020). This highlighted the need for a better molecular and cellular characterization, as well as an improved marker definition of pituitary cells and in particular the pituicytes, ideally leading to the generation of new transgenic tools in rodents and zebrafish. Although β-Ala-Lys-Nε-AMCA was shown to be actively taken up by pituicytes (Anbalagan et al., 2018), we found that other pituitary cells, such as AH progenitors, differentiating hormone-producing cells as well as resident macrophage/microglia were also labeled. We therefore obtained single-cell transcriptomics of the pituicytes along with their neighboring pituitary cells and subsequently characterized their spatial organization in the developing pituitary. In particular, *cyp26b1*+ pituicyte were localized in the dorsal pituitary abutting adenohypophyseal *prop1*+ progenitors.

### Origin of pituicytes and adenohypophyseal cells

Despite the passing of many years since the discovery of pituicytes by Bucy (Bucy, 1930), the exact origin and development of pituicytes remains an enigma. We show here that *cyp26b1+* pituicytes are descendants of *her4.3+* neural plate cells which is in agreement with their radial glia identity evidenced in our scRNA-Seq analysis. This provides the first evidence for the developmental origin of pituicytes prior to formation of the diencephalic infundibulum.

Notably, we found that both the *cyp26b1+* pituicytes and *prop1+* progenitors express many common markers, including the ventral telencephalic radial glia progenitors marker *slc16a9a* (Cosacak et al., 2019). Prop1 is the earliest known exclusive marker of pituitary identity and it is the first pituitary-specific gene in the transcriptional hierarchy (Sornson et al., 1996; Wu et al., 1998). *Prop1*+ cells give rise to all endocrine cells of the AH (Davis et al., 2016). The expression of *slc16a9a* mRNA was not detected in hormone-producing adenohypophyseal cells. However, the long-lived Cherry protein, driven by *slc16a9a* promoter was detected in *pomca*:EGFP+ cells suggesting that it is downregulated during differentiation of *slc16a9a*;*prop1+* progenitor cells into various AH cells. This was further supported by the finding that genetic ablation of *slc16a9a*+ cells leads to a marked reduction in *pomc* and *prl* cells of the AH.

It has been well recognized that the two major pituitary lobes, the AH and NH, do not have the same embryonic origin, with the anterior AH being an oral ectoderm derivative, whereas the posterior NH being derived from neuroectoderm (Alatzoglou et al., 2020; Pearson and Placzek, 2013; Pogoda and Hammerschmidt, 2009; Rizzoti, 2015). Having said that, the anatomical proximity of *prop1+* progenitors and *cyp26b1+* pituicytes and the finding that they shared expression of the ventral forebrain radial glia gene *slc16a9a* inspired us to explore whether they might share early embryonic origins. Using conditional genetic pulse labelling of *her4.3+* neural precursors shortly after the neural plate is formed, we now show that the neural ectoderm gives rise to pituicytes in the NH and a subset of *prop1+* progenitors that likely differentiated into *gh1, prl* and *pomc* cells of the AH. Similarly, labelling of *Sox1*+ progenitors in mouse embryos at early gastrulation stage (6.5dpc) demonstrated contribution of the neural progenitors to both anterior and posterior lobes of the pituitary which give rise to hormone-producing cells of the AH (i.e. LH, GH, TSH and POMC) and Lhx2+ mouse pituicyte respectively. These results challenge the notion that AH cells are exclusively derived from non-neural pre-placodal ectoderm.

Although the pre-placodal ectodermal origin of the AH cells is well established (Couly and Le Douarin, 1988; Dutta et al., 2005; Kouki et al., 2001; Sanchez-Arrones et al., 2017), two studies using radiolabeling in Xenopus or analyzing quail grafted-chick chimeras have suggested that portions of anterior neural ridge (ANR), namely the most anterior neural tissue that meets the non-neural ectoderm, gives rise to at least a part of the adenohypophyseal cells (Couly and Le Douarin, 1985; Eagleson et al., 1986). More recently, lineage tracing of single cells following injection of HRP enzyme to neural plate progenitors at late gastrulation suggested the presence of an intermediate zone comprising multipotential progenitors able to give rise to both the forebrain and the underlying ectoderm, from which RP originates (Cajal et al., 2012).

Although by large these studies support the conclusion of our manuscript, the methods used in these studies lacked single cell resolution. Using more precise (i.e., single cell) genetic lineage tracing and imaging tools, our study provides unequivocal evidence for the contribution of the neural ectoderm to the AH. The concept of multiple embryonic origins of the adenohypophyseal cells is also supported by a recent study demonstrating endodermal contribution to a small subset of AH cells (Fabian et al., 2020).

### Cross-talk between pituicytes and adenohypophyseal progenitors

We have previously shown that pituicytes play an important role in sculpturing the hypothalamo-neurohypophyseal system to adapt its key functions. Pituicyte-derived Slit3, which is locally secreted by pituicytes regulates the maintenance of synaptic neuropeptide levels readily available to be secreted from NH termini (Anbalagan et al., 2019). Pituicyte factors Vegfa, Tgfβ, and Cyp26b1 regulate the formation of a permeable neuroendocrine conduit that bypasses the blood-brain barrier (Anbalagan et al., 2018). The tight anatomical proximity between *cyp26b1+* and *prop1+* cells implied functional cross-talk between pituicytes and adenohypophyseal progenitors.

Thus, we found that genetic and pharmacological perturbations of the pituicytes factor Cyp26b1 affect the transition of *slc16a9a*;*prop1*+ progenitors into differentiated *pomc*+ AH cells. Because Cyp26b1 is a RA degrading enzyme, we attribute the above phenotype to local increase in RA, which acts upon the neighboring *prop1+* progenitors. This is in agreement with other studies, which manifested that Prop1 is required for proper *Aldh1a2* expression and the activation of RA signaling which acts locally on pituitary progenitor cells fated to become hormone-producing cells (Cheung and Camper, 2020; Yoshida et al., 2018). Because *prop1+* cells are intermingled with pituicytes in the developing zebrafish pituitary, we propose that the mutually exclusive expression of *cyp26b1* and *prop1* in intermingled cell populations of the developing zebrafish pituitary ensure tight local regulation of RA signaling at the single cell level. Together with our previous studies, these findings demonstrate that pituicytes, the resident astroglia of the neurohypophysis, orchestrate both functional and developmental properties of the pituitary.

In summary, we present single cell-level molecular composition, cellular cross-talk and lineage relation of the developing zebrafish pituitary. We submit that adenohypophyseal cells are not exclusively derived from the pre-placodal non-neural ectoderm, thereby challenging the prevailing view of the origin of adenohypophyseal cells.

## Materials and Methods

### Animal husbandry

All experiments using zebrafish were approved by the Weizmann Institute’s Institutional Animal Care and Use Committee (Application number #00950121-1 and 01840218-2). The experiments carried out on mice were approved under the UK Animal (scientific procedures) Act (Project license PP8826065). Zebrafish were maintained and bred by standard protocols and according to FELASA guidelines (Aleström et al., 2020). The following transgenes were used in this study, TgBAC(*slc16a9a*:EGFP), TgBAC(*slc16a9a*:GAL4FF), TgBAC(*cyp26b1*:mCherry), Tg(−0.1*pomca*:GFP) (Liu et al., 2003), Tg(*kdrl*:EGFP) (Jin et al., 2005), Tg(UAS:NTR-mCherry) (Davison et al., 2007), Tg(*her4.3*:ERT2CreERT2) (Boniface et al., 2009), Tg(*βactin*:loxP-STOP-loxP-hmgb1-mCherry) (Wang et al., 2011). For larval experiments required imaging, the embryos were kept in the Danieau’s buffer with 0.003% PTU from 24hpf till the day of fixation. *Sox1^CreERT2^*(Sox1^tm3(cre/ERT2)Vep^) (Kicheva et al., 2014) and *Rosa26^RTomato^*(Gt(ROSA)26Sor^tm9(CAG-tdTomato)Hze^) (Madisen et al., 2010) mice were maintained on C57BL/6J background.

### Tol2-mediated Bacterial Artificial Chromosome (BAC) recombineered transgenic zebrafish

BAC transgenes were generated using a Tol2-mediated genomic integration following the protocol as described in (Bussmann and Schulte-Merker, 2011). The BACs containing full zebrafish *slc16a9a* ORF (CH73-306E8) was ordered from BACPAC Resources (CA, UAS). The Tg(*cyp26b1*:mCherry) was generated from BAC clone DKEY-53O14. After insertion of the Tol2 sites into the BAC backbone, a second recombineering step was carried out, replacing the first translated ATG of *slc16a9a* or *cyp26b1* with an eGFP/Gal4FF or mCherry reporter cassette. The targeting products containing the iTol2-cmlc2:mTurq cassette with m13 tags, eGFP/Gal4FF or mCherry cassettes with *slc16a9a*/*cyp26b1* homology arms for recombineering were amplified by PCR. 200ng/µl of the final BAC DNAs (*slc16a9a*:eGFP/*slc16a9a*:Gal4FF) were then injected into 1-2 cells stage AB lines with 25ng/µl transposase mRNA. Embryos showing fluorescence in the heart were raised to adulthood and then mated to select for successful germline transmission. Integration of the targeting products was assessed using the control primer pairs. All primers used are listed in Table 3. At least two founders were selected for each transgenic line and outcrossed with AB for maintenance.

### Neurohypophyseal cell enrichment

Neurohypophyseal cells were enriched by intraperitoneally injecting 5µl of 4.6mM β-Ala-Lys-Nε-AMCA (Biotrend #BP0352) into 37-day-old juvenile, or 10µl into 5 or 7 months-old male Tg(*pomca*:EGFP). After 3 hours, the pituitaries of each age group were dissected and dispersed into single cell suspensions as described in (Anbalagan et al., 2018) with slight modifications. Briefly, 5 pituitaries from each age group were dissected and pooled in a 1.5ml tube containing 1ml ice-cold HEPES buffered saline (HBS), which was later changed to 250μl ice-cold phosphate buffered saline (PBS)+/+ with prewarmed Liberase TM (Roche, Switzerland) for 12 minutes at 30 degrees C. The tissues were then further dissociated with 750μl of TrypLE Express (Thermo Fisher, USA) and 16μl DNaseI (Qiagen, Germany) for 6 minutes at room temperature and stopped with 50μl FBS. Dissociated cells were pelleted at 500g for 5 minutes at 4°C, thereafter resuspended in 500μl ice-cold resuspension buffer (Leibovitz L-15 with 0.3mM Glutamine (GIBCO, Thermo Fisher, USA), penicillin 50 U/mL + streptomycin 0.05 mg/mL, FBS 1%, BSA 0.04%) and passed through a 40μm strainer (BD, USA) to obtain single cell suspension. Before FACS sorting, propidium iodide was added to distinguish dead cells. FACS was conducted with a 100μm nozzle from either SORP-FACSAriaII (BD) or FACSAriaIII (BD, USA) system. The cells were first gated by SSC-A and FSC-A strategy, live cells were selected by negative staining for propidium iodide, singlets were then gated using FSC-H and FSC-A method. Single AMCA-positive, eGFP-negative cell was sorted into each well of a 384-well plate containing lysis buffer with barcoded poly(T) reverse-transcription (RT) primers (Jaitin et al., 2014). Four wells were left empty as negative controls. The plates were then centrifugated and snap-frozen on dry ice then stored at −80°C prior to single-cell RNA sequencing library preparation. For each age group, two 384-well plates of cells were collected from 10-32 pituitaries (Supplementary Table 1).

### Mars-Seq single-cell RNA sequencing

The Mars-Seq single-cell library was prepared according to (Jaitin et al., 2014). In short, mRNA from the single cell was barcoded and converted to cDNA in the 384-well plate, then pooled and *in vitro* transcribed to RNA before fragmented into a library. The library was then ligated, reverse transcribed, and PCR amplified to attach the pool barcodes and Illumina sequencing codes. The library quality and concentration for each pool were checked before sequencing using an Illumina NextSeq 500 machine (Illumina, USA) at a median sequencing depth around 40,000 reads per cell.

### Data and software availability

The zebrafish neurohypophysis single-cell transcriptome in this study is available in Gene Expression Omnibus (GEO) (www.ncbi.nlm.nih.gov/geo) with accession number: GSE203075.

### Bioinformatic analyses

Sequences with low quality RMT were filtered out, pool-barcode and well-barcode-RMT were extracted to create fastq files, and then mapped to the zebrafish genome (danRer10). Single cell analysis was performed using the Seurat package v 3.1.5 (Butler et al., 2018). Cells with >30% mitochondria gene expression as well as cells with extreme top or bottom UMI expression were removed leaving a total of 1281 cells from all samples. Expression values were normalized and scaled using the functions NormalizeData (normalization.method=“LogNormalize”, scale.factor= 10000) and ScaleData (vars.to.regress = c(“nCount_RNA”, “percent.mito”). Clusters were determined using FindNeighbors (dims=1:6) and FindClusters (reduction.type = “pca”, resolution = 0.6). Single cell reanalysis for Fabian et al. was conducted in the same pipeline as above. Clusters were determined using FindNeighbors (dims=1:20) and FindClusters (reduction.type = “pca”, resolution = 1.2).

The over representation analysis (ORA) (Boyle et al., 2004) was performed as described in our previous publication (Chen Q et al., 2020) using an in house developed R script. ORA was done also using cluster cell markers of scRNA from the adult mouse NH scRNA-Seq data, and the human fetus pituitary scRNA-Seq data (Chen Q et al., 2020; Franzen et al., 2019; Zhang et al., 2020). All scRNA-Seq markers were filtered by an average fold change >=1.5 and adjusted pvalue<=0.05. The hypergeometric *p*-value for ORA was calculated by R (Kachitvichyanukul and Schmeiser, 1985), and the FDR was achieved by adjusting the *p-*value per cluster using Benjamini and Hochberg (Benjamini and Hochberg, 1995). The data were further filtered by enrichment FDR per cluster<=0.05, with the number and symbols of genes in each data set summarized in Table 2.

### In situ hybridization and immunostaining in zebrafish

Whole-mount *in situ* hybridization was conducted as described in (Borodovsky et al., 2009; Schulte-Merker, 2002). RNA probes were synthesized by amplifying a partial coding sequence of the genes with a T7 tagged reversed primer. The PCR products were then purified and reverse transcribed with digoxigenin labeling using a DIG RNA labelling kit (Roche, Switzerland). For dissected adult pituitary or 5-8 dpf larvae, 50 minutes or 30-40 minutes of proteinase K digestion was conducted.

Hybridization chain reaction (HCR) *in situ* hybridization v3.0 probes were custom designed and synthesized by Molecular Instruments (USA). Whole-mount HCR *in situ* hybridization of zebrafish larvae was performed according to (Choi et al., 2016, 2018) with slight modifications: the final amount of probes during incubation was increased to 5pmol for each set, and the samples were fixed in 4% paraformaldehyde (PFA) after detection stage and washed in 0.1% Tween20 PBS 3 times before proceeding with amplification stage.

Antibody staining for eGFP (GFP Rabbit Polyclonal #A11122 or GFP Chicken Polyclonal #A10262, Invitrogen, Thermo Fisher, USA) and/or mCherry (Living Colors DsRed Polyclonal Antibody #632496, Takara, Japan) after *in situ* hybridization were carried out as described in (Chen Q et al., 2020). The samples were then mounted in 75% glycerol, and imaged using a Zeiss LSM800 confocal microscope with 40X oil lens. Maximum intensity projections and further imaging quantifications were conducted with Fiji-ImageJ (Fiji, Japan). Video of the 3D images was made by Imaris v9.5.0 (Oxford Instruments, UK).

### Processing and immunostaining of mouse tissues

Embryos genotyping was performed by examining live fluorescence. For late gestation embryos, positive heads were dissected, and the pituitary exposed. Fixation of dissected heads or 11.5dpc embryos was performed overnight at 4°C in 4% PFA. Samples were cryoprotected in sucrose and embedded in OCT for cryosection. Immunofluorescence were performed using Rabbit anti-GH, LH, and POMC, Guinea Pig anti-TSH (all anti hormone antibodies originate from the NHPP), Goat anti-SOX2 (Immune System, GT15098) and Rabbit anti-LHX2 (Abcam, ab184337). Anti-rabbit, anti-guinea-pig or anti-goat secondary antibodies conjugated to Alexa-Fluor 488, −568, or −647 were used for immunolabelling. Imaging was performed using a Leica SPE confocal microscope.

### Lineage tracing

The transgenic zebrafish lines *her4.3*:ERT2CreERT2 (Boniface et al., 2009) and *βactin*:loxP-STOP-loxP-hmgb1-mCherry (Wang et al., 2011) were kindly provided by Dr. Laure Bally-Cuif (Pasteur Institute, Paris). For CRE-dependent genetic labeling of *her4.3+* precursors, zebrafish embryos were dechorionated with 1mg/ml pronase at 9 hpf and washed thoroughly with fresh Danieau’s buffer before being treated with 10μM (Z)-4-hydroxytamoxifen (#3412, Tocris, UK) for 2h at 28°C in the dark. At the end of the treatment, the embryos were washed multiple times with fresh Danieau’s buffer and transferred to a new plate that was placed in an incubator at 28°C. They were then visually inspected for mCherry expression at 3dpf under a fluorescent stereomicroscope, and fixed at 6dpf in 4% PFA at 4°C overnight. The larvae used for either immunostaining or HCR *in situ* hybridization with immunostaining were kept in absolute methanol at −20°C overnight as described in the previous session. Two control groups were also imaged to assess the leakiness of the recombineering procedure as well as and tissue toxicity. These were the 0.1% ethanol treated ERT2CreERT2 and loxP-STOP-loxP-hmgb1-mCherry double positive siblings, and 4-OHT treated siblings without ERT2CreERT2/loxP-STOP-loxP-hmgb1-mCherry. For the lineage tracing of mouse SOX1+ progenitors, homozygous *Rosa26^RTomato^* female mice were mated with *Sox1^CreERT2/+^* studs to produce experimental embryos. Day of vaginal plug was counted as 0.5dpc. Cre activity was induced by a single tamoxifen treatment (0.2mg/g body weight) in pregnant females.

### Genetic Cell ablation

For nitroreductase-mediated conditional cell ablation, 3dpf embryos harboring TgBAC(*slc16a9a*:GAL4FF;UAS:NTR-mCherry) transgene and their respective controls were treated with 5μM nifuripirinol (NFP) for 24h at 28°C. After the treatment, and the larvae were washed 3 times (10min each) with fresh Danieau’s buffer, thereafter transferred to a new plate and maintained at 28°C for additional 48h to recover. Larvae were then fixed at 6dpf in 4% PFA at 4°C overnight. The larvae were either mounted in 75% glycerol and subjected to imaging or kept in absolute methanol at −20°C for further HCR *in situ* and immunostaining as described in the previous session. 0.1% DMSO treated siblings, and NFP treated Gal4FF/UAS negative siblings were also imaged and quantified as controls. Identical imaging parameters were used for all groups from each cohort.

### R115866 (Talarozole) treatment

Embryos were treated for 24h with 2μM of R115866/Talarozole (MedChemExpress, USA) at 28°C between 4 to 5dpf and fixed in 4% PFA at 4°C overnight. The larvae were either mounted in 75% glycerol for imaging or kept in absolute methanol at −20°C overnight for whole-mount *in situ* hybridization and antibody staining as described in the previous section. Controls were mock-treated with 0.1% DMSO. In each experiment, the imaging parameters were kept the same for all groups.

### Image quantifications and statistical analyses

Normalized mean intensity analysis of the z-stack image was developed in our lab using ImageJ. The target channel and range of z-stacks were selected, then zproject function was run in projection type ‘*sum slices’*. Mean intensities of two regions of interest (ROIs) were measured: one for the major image and another for the background control. Areas of the ROIs were kept the same for each group within one cohort. The normalized mean intensity = mean intensity _major ROI_ – mean intensity _background ROI_. The percentage of reduced signals = (group mean intensity _DMSO_– group mean intensity _treatment_)/group mean intensity _DMSO_ *100%. Colocalization analysis was conducted using an ImageJ plugin: stack colocalization analyzer (developed by Alex Herbert, University of Sussex) with the following parameters: method Otsu, subtract threshold, permutations 50, minimum shift 9, maximum shift 16, and significance 0.05. Images without whole-mount *in situ* and immunostaining were proceeded directly; images with whole-mount *in situ* and antibody staining were first batch cropped using an ImageJ macro and then batch analyzed the colocalization using an imageJ macro.

Graphs were generated with Graphpad Prism 6 (CA, UAS). Data from all groups were checked for normality by D’Agostino & Pearson omnibus, Shapiro-Wilk, and KS normality tests. Based on the normality, the significance of the three groups comparisons were calculated by either Kruskal-Wallis test or ordinary one-way ANOVA with the posteriori Dunn’s multiple comparison test to the control; the significance of the two groups comparison were calculated by unpaired student’s t-test by either Welch’s correction or Mann-Whitney test.

## Acknowledgements

We thank Roy Hofi, Estar Regev and fish facility personnel; Dr. Shuang-Yin Wang, Gil Stelzer Dr. Ron Rotkopf, and Tal Fisher for helping with bioinformatic and image analyses; Dr. Isabelle Foucher and Dr. Laure Bally-Cuif for offering the *her4.3* transgenic lines. Genia Brodsky for the figure graphics. G.L. lab is supported by the Israel Science Foundation (#349/21); US-Israel Bi-National Science Foundation (#2017325); Israel Ministry of Science (#3-16548); Hedda, Alberto, and David Milman Baron Center for Research on the Development of Neural Networks; Sagol Institute for Longevity Research; Maurice and Vivienne Wohl Biology Endowment and Foundation for Higher Education and Culture (FFHEC). K.R. was supported by the Francis Crick Institute which receives its core funding from Cancer Research UK (FC001107), the UK Medical Research Council (FC001107), and the Wellcome Trust (FC001107). Q.C. was supported by the European Zebrafish Society Short Term Scientific Mission (STSM) Fellowship. G.L. is an incumbent of the Elias Sourasky Professorial Chair.

---

**Supplementary Figure 1.**
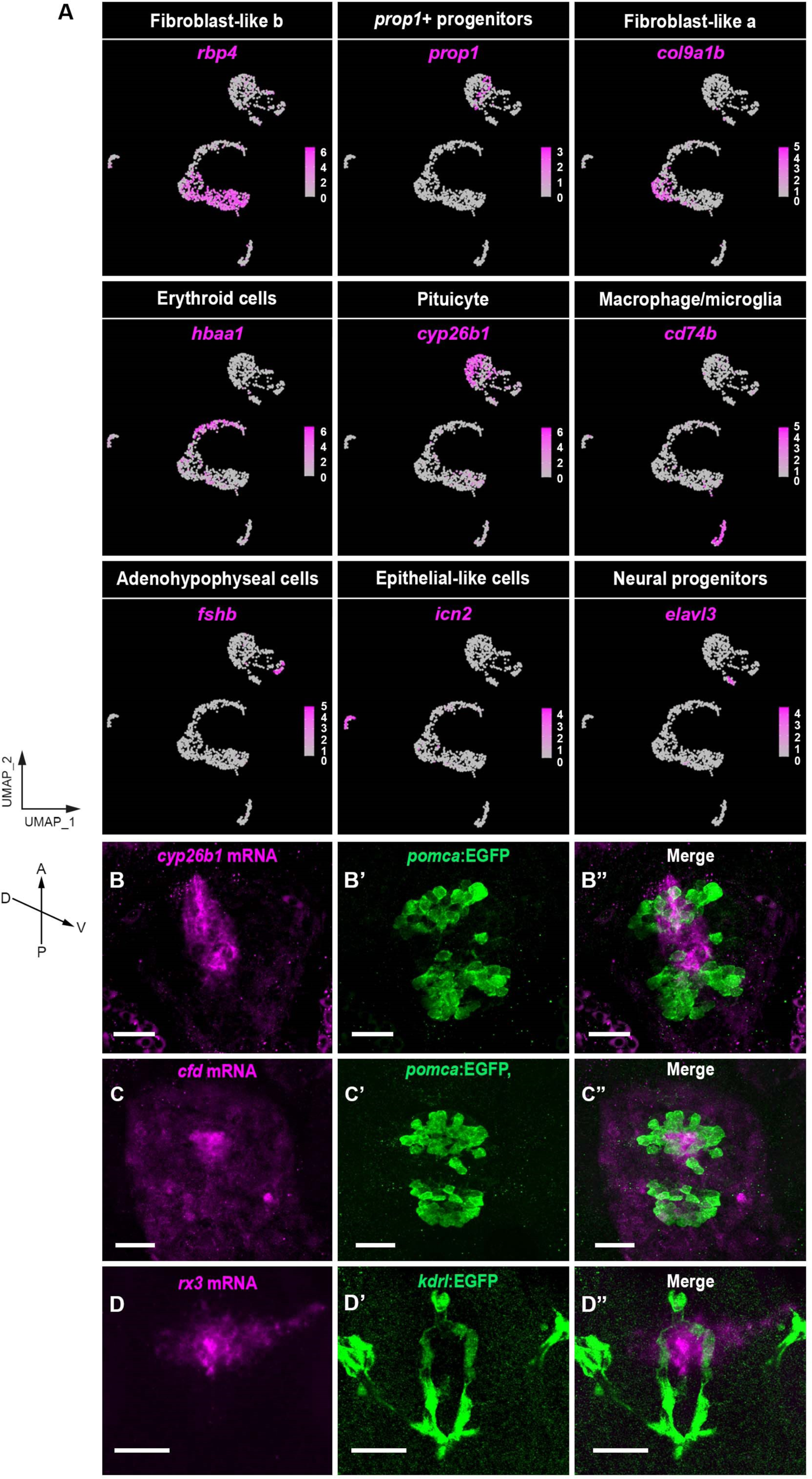
Cell-type specific feature genes and validations of selected pituicyte markers in larval zebrafish. (**A**) Featured plots in which specific marker genes representing each cluster are superimposed on the UMAP. The cluster names and marker genes are indicated at the top of each panel. The gene expression levels are color-coded with high expression in magenta and low expression in gray. (**B-D**) Transgenic 5-8 day-old larvae harboring either the adenohypophyseal reporter [Tg(*pomca*:EGFP)**(B, C)**] or the vascular endothelial reporter [Tg(*kdrl*:EGFP) **(D)**] were used as anatomical landmarks to indicate the location of pituitary by adenohypophyseal Pomc cells or the hypophyseal vasculature. Larvae were subjected to whole-mount *in situ* hybridization using RNA probes directed to specific pituicytes markers revealed by scRNA-Seq. Maximum intensity projected confocal images show the pituicyte markers *cyp26b1* (**B**), *cfd (***C**), and *rx3* (**D**). C, caudal; D, dorsal; R, rostral; V, ventral. Scale bars, 20μm.

**Supplementary Figure 2.**
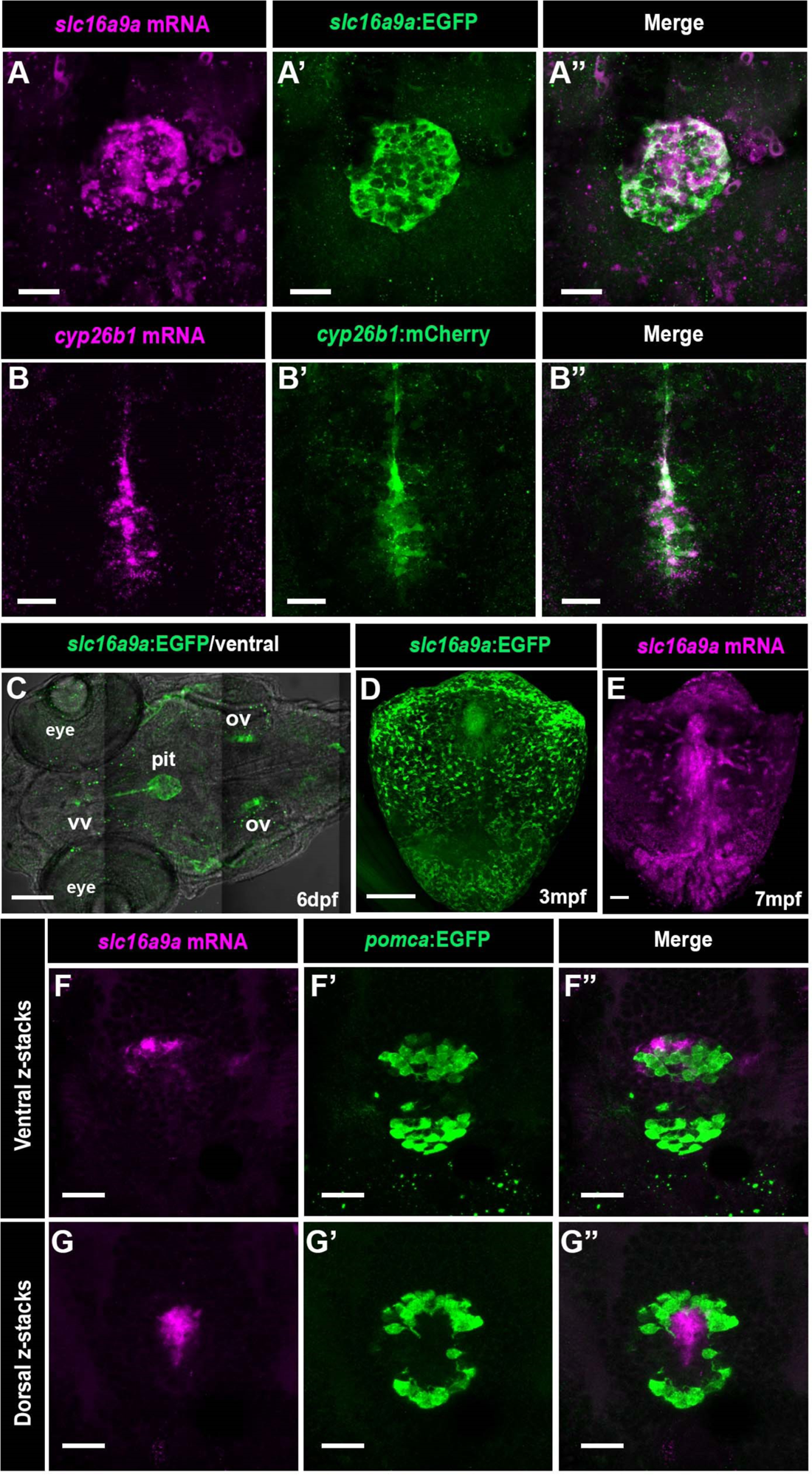
Tg(*slc16a9a*:EGFP) and Tg(*cyp26b1*:mCherry) reporters faithfully represent endogenous mRNA expression in the pituitary. (**A-B**) Maximal projected Z-stack confocal images of the pituitary region from an 8dpf transgenic Tg(*slc16a9a*:EGFP) (**A**) or a 6dpf Tg(*cyp26b1*:mCherry) larvae (**B**) which were subjected to *in situ* hybridization of mRNA probes directed to *slc16a9a or cyp26b1*, respectively, followed by anti-EGFP/anti-mCherry immunostaining. Scale bars, 20μm. (C) Confocal Z-stack image of a 6dpf transgenic Tg(*slc16a9a*:EGFP) larva, which was immunostained with anti-EGFP antibody and overlayed with brightfield image (ventral view) showing EGFP signals in the pituitary, the ventral nucleus of the ventral telencephalic area (vv), and otic vesicle (ov). pit, pituitary. Scale bar, 100μm. (D) Dissected pituitary from a 3 months-old transgenic Tg(*slc16a9a*:EGFP) adult, which was immunostained with an anti-EGFP antibody. Scale bar, 100μm. (E) *In situ* hybridization of *slc16a9a* mRNA expression in a dissected pituitary of a 7 months-old adult. Scale bar, 50μm. (**F,G**) Ventral (**F-F”**) and dorsal (**G-G”**)serial confocal Z-stacks of a 8dpf Tg(*pomca*:EGFP) larva, which was subjected to *slc16a9a* mRNA *in situ* hybridization followed by immunostaining with an anti-EGFP antibody. Scale bars, 20μm.

**Supplementary Figure 3.**
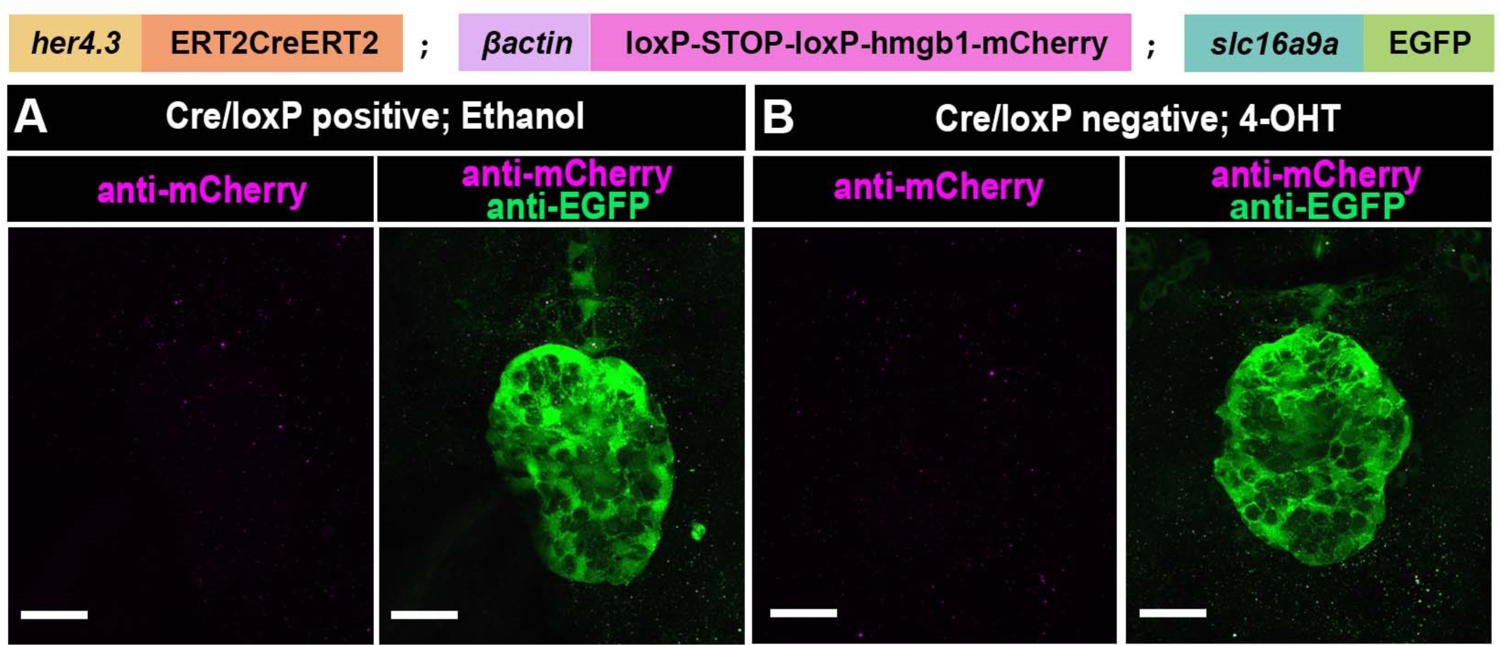
*her4.3*:ERT2CreERT2 driver is tightly controlled by 4-OHT. (A) Triple transgenic embryo Tg(*her4.3*:ERT2CreERT2;*βactin*:loxP-STOP-loxP-hmgb1-mCherry;*slc16a9a*:EGFP) were genotyped before treatment with 0.1% ethanol between 9-11hpf, and co-immunostained with anti-mCherry and anti-EGFP staining at 6dpf. Scale bar, 20μm. (B) *slc16a9a*:EGFP-positive;Cre/LoxP-negative embryos derived from a triple transgenic cross shown in **A** were treated with 10μM 4-OHT followed by anti-mCherry and anti-EGFP immunostaining. Scale bar, 20μm.

**Supplementary Figure 4.**
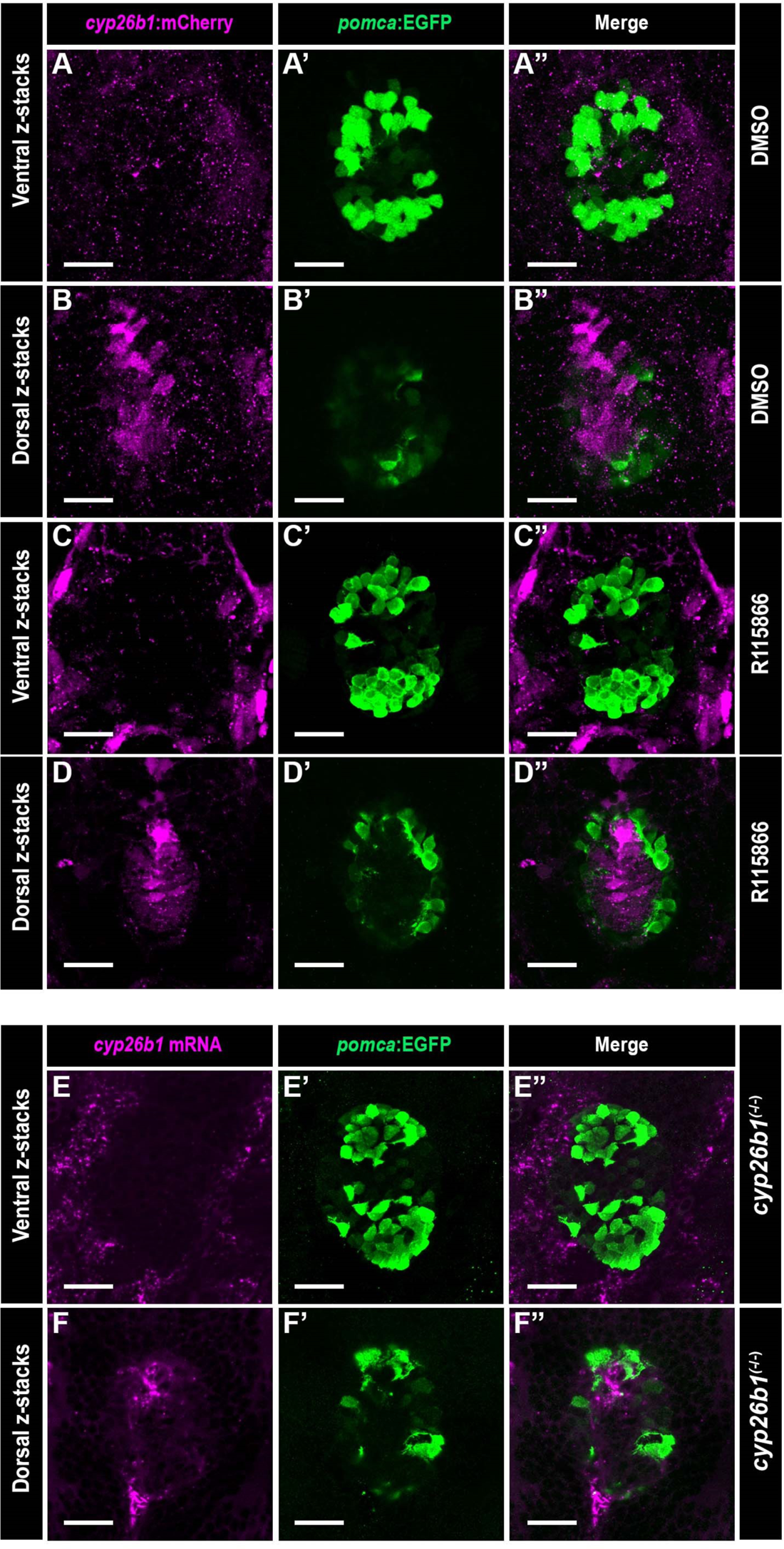
*cyp26b1* mRNA and *cyp26b1:*mCherry reporter expression do not overlap with and Pomc cells. (**A-D**) Confocal images showing either ventral (**A, C**) or dorsal (**B, D**) serial Z-stack of the pituitary in 5dpf Tg(*cyp26b1*:mCherry;*pomca*:EGFP) larvae that were treated with either 0.1% DMSO (n=9) or 2μM R115866 (n=10) between 4-5 days and immunostained with anti-mCherry and anti-EGFP antibodies. No overlapping mCherry/EGFP signals were detected in any of the samples. Scale bars, 20μm. (**E-F**) Confocal images showing either ventral (**E**) or dorsal (**F**) serial Z-stack of *cyp26b1* mRNA and Tg(*pomca*:EGFP) reporter in the pituitary of a 5dpf *cyp26b1*^(-/-)^ (n=8). No overlapping *cyp26b1*/EGFP signals were detected. Scale bars, 20μm.

**Movie 1.** Sequential Z-stack slices showing the 3D structure in a 6dpf Tg(*slc16a9a*:EGFP); Tg(*cyp26b1*:mCherry) pituitary with *prop1* mRNA, anti-mCherry, and anti-EGFP staining. Scale bar, 10μm.

**Table 1.** Sample information of β-Ala-Lys-Nε-AMCA enriched neurohypophyseal cells collected for Mars-Seq scRNA-Seq and normalized differentially expressed genes in each cluster filtered by average fold change >=2.0 and adjusted p-value <= 0.05.

**Table 2.** ORA comparing differentially expressed genes from all clusters in this study to that from the mouse neurohypophyseal scRNA-Seq and human fetus pituitary scRNA-Seq. Gene lists for all species were filtered by average fold change >=1.5 and adjusted *p*-value <=0.05. Data from ORA were further filtered by enrichment FDR per cluster <= 0.05.

**Table 3.** Oligonucleotides used in this study.

## Reference

1. Agarwal, G., Bhatia, V., Cook, S., and Thomas, P.Q. (2000). Adrenocorticotropin deficiency in combined pituitary hormone deficiency patients homozygous for a novel PROP1 deletion. J. Clin. Endocrinol. Metab. 85, 4556–4561.

2. Alatzoglou, K.S., Gregory, L.C., and Dattani, M.T. (2020). Development of the Pituitary Gland. Compr. Physiol. 10, 389–413.

3. Aleström, P., D’Angelo, L., Midtlyng, P.J., Schorderet, D.F., Schulte-Merker, S., Sohm, F., and Warner, S. (2020). Zebrafish: Housing and husbandry recommendations. Lab. Anim. 54, 213– 224.

4. Anbalagan, S., Gordon, L., Blechman, J., Matsuoka, R.L., Rajamannar, P., Wircer, E., Biran, J., Reuveny, A., Leshkowitz, D., Stainier, D.Y.R., et al. (2018). Pituicyte cues regulate the development of permeable neuro-vascular interfaces. Dev. Cell 47, 711–726.

5. Anbalagan, S., Blechman, J., Gliksberg, M., Gordon, L., Rotkopf, R., Dadosh, T., Shimoni, E., and Levkowitz, G. (2019). Robo2 regulates synaptic oxytocin content by affecting actin dynamics. Elife 8, e45650.

6. Angotzi, A.R., Mungpakdee, S., Stefansson, S., Male, R., and Chourrout, D. (2011). Involvement of Prop1 homeobox gene in the early development of fish pituitary gland. Gen. Comp. Endocrinol. 171, 332–340.

7. De Beer, G.R. (1924). The Evolution of the Pituitary. J. Exp. Biol. 1, 271–291.

8. Benjamini, Y., and Hochberg, Y. (1995). Controlling the False Discovery Rate: A Practical and Powerful Approach to Multiple Testing. J. R. Stat. Soc. Ser. B 57, 289–300.

9. Bergemann, D., Massoz, L., Bourdouxhe, J., Carril Pardo, C.A., Voz, M.L., Peers, B., and Manfroid, I. (2018). Nifurpirinol: A more potent and reliable substrate compared to metronidazole for nitroreductase-mediated cell ablations. Wound Repair Regen. Off. Publ. Wound Heal. Soc. [and] Eur. Tissue Repair Soc. 26, 238–244.

10. Biran, J., Blechman, J., Wircer, E., and Levkowitz, G. (2018). Development and Function of the Zebrafish Neuroendocrine System. Model Anim. Neuroendocrinol. 101–131.

11. Boniface, E.J., Lu, J., Victoroff, T., Zhu, M., and Chen, W. (2009). FlEx-based transgenic reporter lines for visualization of Cre and Flp activity in live zebrafish. Genesis 47, 484–491.

12. Borodovsky, N., Ponomaryov, T., Frenkel, S., and Levkowitz, G. (2009). Neural protein Olig2 acts upstream of the transcriptional regulator Sim1 to specify diencephalic dopaminergic neurons. Dev. Dyn. 238, 826–834.

13. Boyle, E.I., Weng, S., Gollub, J., Jin, H., Botstein, D., Cherry, J.M., and Sherlock, G. (2004). GO::TermFinder - Open source software for accessing Gene Ontology information and finding significantly enriched Gene Ontology terms associated with a list of genes. Bioinformatics 20, 3710–3715.

14. Bucy, P.C. (1930). The pas nervosa of the bovine hypophysis. J. Comp. Neurol. 50, 505–519.

15. Bussmann, J., and Schulte-Merker, S. (2011). Rapid BAC selection for tol2-mediated transgenesis in zebrafish. Development 138, 4327–4332.

16. Butler, A., Hoffman, P., Smibert, P., Papalexi, E., and Satija, R. (2018). Integrating single-cell transcriptomic data across different conditions, technologies, and species. Nat. Biotechnol. 36, 411–420.

17. Cajal, M., Lawson, K.A., Hill, B., Moreau, A., Rao, J., Ross, A., Collignon, J., and Camus, A. (2012). Clonal and molecular analysis of the prospective anterior neural boundary in the mouse embryo. Development 139, 423–436.

18. Chen Q., Leshkowitz, D., Blechman, J., and Levkowitz, G. (2020). Single-cell molecular and cellular architecture of the mouse neurohypophysis. Eneuro ENEURO.0345–19.2019.

19. Cheung, L.Y.M., and Camper, S.A. (2020). PROP1-Dependent Retinoic Acid Signaling Regulates Developmental Pituitary Morphogenesis and Hormone Expression. Endocrinology 161.

20. Cheung, L.Y.M., George, A.S., McGee, S.R., Daly, A.Z., Brinkmeier, M.L., Ellsworth, B.S., and Camper, S.A. (2018). Single-cell RNA sequencing reveals novel markers of male pituitary stem cells and hormone-producing cell-types. Endocrinology 159, 3910–3924.

21. Choi, H.M.T., Calvert, C.R., Husain, N., Huss, D., Barsi, J.C., Deverman, B.E., Hunter, R.C., Kato, M., Lee, S.M., Abelin, A.C.T., et al. (2016). Mapping a multiplexed zoo of mRNA expression. Development 143, 3632–3637.

22. Choi, H.M.T., Schwarzkopf, M., Fornace, M.E., Acharya, A., Artavanis, G., Stegmaier, J., Cunha, A., and Pierce, N.A. (2018). Third-generation in situ hybridization chain reaction: multiplexed, quantitative, sensitive, versatile, robust. Development 145.

23. Clasadonte, J., and Prevot, V. (2018). The special relationship: Glia-neuron interactions in the neuroendocrine hypothalamus. Nat. Rev. Endocrinol. 14, 25–44.

24. Cosacak, M.I., Bhattarai, P., Reinhardt, S., Petzold, A., Dahl, A., Zhang, Y., and Kizil, C. (2019). Single-Cell Transcriptomics Analyses of Neural Stem Cell Heterogeneity and Contextual Plasticity in a Zebrafish Brain Model of Amyloid Toxicity. Cell Rep. 27, 1307–1318.e3.

25. Couly, G., and Le Douarin, N.M. (1988). The fate map of the cephalic neural primordium at the presomitic to the 3-somite stage in the avian embryo. Development 103, 101–113.

26. Couly, G.F., and Le Douarin, N.M. (1985). Mapping of the early neural primordium in quail-chick chimeras. I. Developmental relationships between placodes, facial ectoderm, and prosencephalon. Dev. Biol. 110, 422–439.

27. Couly, G.F., and Le Douarin, N.M. (1987). Mapping of the early neural primordium in quail-chick chimeras. II. The prosencephalic neural plate and neural folds: implications for the genesis of cephalic human congenital abnormalities. Dev. Biol. 120, 198–214.

28. Curado, S., Stainier, D.Y.R., and Anderson, R.M. (2008). Nitroreductase-mediated cell/tissue ablation in zebrafish: a spatially and temporally controlled ablation method with applications in developmental and regeneration studies. Nat. Protoc. 3, 948–954.

29. Davis, S.W., Ellsworth, B.S., Peréz Millan, M.I., Gergics, P., Schade, V., Foyouzi, N., Brinkmeier, M.L., Mortensen, A.H., and Camper, S.A. (2013). Pituitary gland development and disease: from stem cell to hormone production. Curr. Top. Dev. Biol. 106, 1–47.

30. Davis, S.W., Keisler, J.L., Pérez-Millán, M.I., Schade, V., and Camper, S.A. (2016). All Hormone-Producing Cell Types of the Pituitary Intermediate and Anterior Lobes Derive from Prop1-Expressing Progenitors. Endocrinology 157, 1385–1396.

31. Davison, J.M., Akitake, C.M., Goll, M.G., Rhee, J.M., Gosse, N., Baier, H., Halpern, M.E., Leach, S.D., and Parsons, M.J. (2007). Transactivation from Gal4-VP16 transgenic insertions for tissue-specific cell labeling and ablation in zebrafish. Dev. Biol. 304, 811–824.

32. Dellmann, H.D., and Sikora, K. (1981). Pituicyte fine structure in the developing neural lobe of the rat. Dev. Neurosci. 4, 89–97.

33. Dutta, S., Dietrich, J.-E., Aspöck, G., Burdine, R.D., Schier, A., Westerfield, M., and Varga, Z.M. (2005). pitx3 defines an equivalence domain for lens and anterior pituitary placode. Development 132, 1579–1590.

34. Eagleson, G.W., Jenks, B.G., and Van Overbeeke, A.P. (1986). The pituitary adrenocorticotropes originate from neural ridge tissue in Xenopus laevis. J. Embryol. Exp. Morphol. 95, 1–14.

35. Fabian, P., Tseng, K.-C., Smeeton, J., Joseph, L.J., Dong, P.D.S., Cerny, R., and Crump, G. (2020). Lineage analysis reveals an endodermal contribution to the vertebrate pituitary. Science. 370 (6515), 463–467.

36. Fletcher, P.A. (2021). Transcriptomic heterogeneity of Sox2-expressing pituitary cells. BioRxiv Neurosci. 1–28.

37. Franzen, O., Gan, L., and Björkegren, J.L.M. (2019). PanglaoDB: a web server for exploration of mouse and human single-cell RNA. Database 2019, 1–9.

38. Gutnick, A., and Levkowitz, G. (2012). The Neurohypophysis: Fishing for New Insights. J. Neuroendocrinol. 24, 973–974.

39. Hatton, G.I. (1988). Pituicytes, glia and control of terminal secretion. J. Exp. Biol. 139, 67–79.

40. Herring, P.T. (1908). Histological Appearance of Mammalian Pituitary. Quart. Journ. Exp. Phys 1.

41. Imoesi, P.I., Bowman, E.E., Stoney, P.N., Matz, S., and McCaffery, P. (2019). Rapid Action of Retinoic Acid on the Hypothalamic Pituitary Adrenal Axis. Front. Mol. Neurosci. 12.

42. Jaitin, D.A., Kenigsberg, E., Keren-Shaul, H., Elefant, N., Paul, F., Zaretsky, I., Mildner, A., Cohen, N., Jung, S., Tanay, A., et al. (2014). Massively Parallel Single-Cell RNA-Seq for Marker-Free Decomposition of Tissues into Cell Types. Science. 343 (6172), 776–779.

43. Jin, S.-W., Beis, D., Mitchell, T., Chen, J.-N., and Stainier, D.Y.R. (2005). Cellular and molecular analyses of vascular tube and lumen formation in zebrafish. Development 132, 5199–5209.

44. Kachitvichyanukul, V., and Schmeiser, B. (1985). Computer generation of hypergeometric random variates. J. Stat. Comput. Simul. 22, 127–145.

45. Kawamoto, K., and Kawashima, S. (1984). Ultrastructural changes and proliferation of pituicytes in mouse posterior lobe during water deprivation and rehydration. Acta Anat. (Basel). 119, 136–141.

46. Kicheva, A., Bollenbach, T., Ribeiro, A., Valle, H.P., Lovell-Badge, R., Episkopou, V., and Briscoe, J. (2014). Coordination of progenitor specification and growth in mouse and chick spinal cord. Science 345, 1254927.

47. Kouki, T., Imai, H., Aoto, K., Eto, K., Shioda, S., Kawamura, K., and Kikuyama, S. (2001). Developmental origin of the rat adenohypophysis prior to the formation of Rathke’s pouch. Development 128, 959–963.

48. Larkin, S., and Ansorge, O. (2017). Development and microscopic anatomy of the pituitary gland. In Endotext [Internet], L. De Groot, G. Chrousos, and K. Dungan, eds. (South Dartmouth (MA): MDText.com, Inc.), p. 2000.

49. Leng, G., Pineda, R., Sabatier, N., and Ludwig, M. (2015). 60 YEARS OF NEUROENDOCRINOLOGY: The posterior pituitary, from Geoffrey Harris to our present understanding. J. Endocrinol. 226, T173–85.

50. Liu, N.-A., Huang, H., Yang, Z., Herzog, W., Hammerschmidt, M., Lin, S., and Melmed, S. (2003). Pituitary Corticotroph Ontogeny and Regulation in Transgenic Zebrafish. Mol. Endocrinol. 17, 959–966.

51. Madisen, L., Zwingman, T.A., Sunkin, S.M., Oh, S.W., Zariwala, H.A., Gu, H., Ng, L.L., Palmiter, R.D., Hawrylycz, M.J., Jones, A.R., et al. (2010). A robust and high-throughput Cre reporting and characterization system for the whole mouse brain. Nat. Neurosci. 13, 133–140.

52. Marin, F., Boya, J., and Lopez-Carbonell, A. (1989). Immunocytochemical localization of vimentin in the posterior lobe of the cat, rabbit and rat pituitary glands. Acta Anat. (Basel). 134, 184–190.

53. Mayran, A., Sochodolsky, K., Khetchoumian, K., Harris, J., Gauthier, Y., Bemmo, A., Balsalobre, A., and Drouin, J. (2019). Pioneer and nonpioneer factor cooperation drives lineage specific chromatin opening. Nat. Commun. 10, 3807.

54. Menuet, A., Pellegrini, E., Brion, F., Gueguen, M.-M., Anglade, I., Pakdel, F., and Kah, O. (2005). Expression and estrogen-dependent regulation of the zebrafish brain aromatase gene. J. Comp. Neurol. 485, 304–320.

55. Miyata, S. (2017). Advances in understanding of structural reorganization in the hypothalamic neurosecretory system. Front. Endocrinol. (Lausanne). 8, 1–18.

56. Murphy, D., Konopacka, A., Hindmarch, C., Paton, J.F.., Sweedler, J. V, Gillette, M.U., Ueta, Y., Grinevich, V., Lozic, M., and Japundzic-Zigon, N. (2012). The hypothalamo-neurohypophyseal system: from genome to physiology. J. Endocrinol. 24, 539–553.

57. Nelson, C.H., and Isoherranen, B.R.B. and N. (2013). Therapeutic Potential of the Inhibition of the Retinoic Acid Hydroxylases CYP26A1 and CYP26B1 by Xenobiotics. Curr. Top. Med. Chem. 13, 1402–1428.

58. Pearson, C.A., and Placzek, M. (2013). Development of the Medial Hypothalamus: Forming a Functional Hypothalamic-Neurohypophyseal Interface. Curr. Top. Dev. Biol. 106, 49–88.

59. Pevny, L.H., Sockanathan, S., Placzek, M., and Lovell-Badge, R. (1998). A role for SOX1 in neural determination. Development 125, 1967–1978.

60. Pogoda, H.-M., and Hammerschmidt, M. (2009). How to make a teleost adenohypophysis: molecular pathways of pituitary development in zebrafish. Mol. Cell. Endocrinol. 312, 2–13.

61. Rathke, H. (1838). Ueber die Entstehung der Glandula pituitaria. Muller’s Arch. S. 482.

62. Rizzoti, K. (2015). Genetic regulation of murine pituitary development. J. Mol. Endocrinol. 54, R55–73.

63. Robinson, A.G., and Verbalis, J.G. (2003). The posterior pituitary gland. In Williams Textbook of Endocrinology, P. Larson, H. Kronengerg, S. Melmed, and K. Polonsky, eds. (Philadelphia: WB Saunders), pp. 281–330.

64. Rodríguez, E., Guerra, M., Peruzzo, B., and Blázquez, J.L. (2019). Tanycytes: A rich morphological history to underpin future molecular and physiological investigations. J. Neuroendocrinol. 31, 0–2.

65. Sanchez-Arrones, L., Sandonís, Á., Cardozo, M.J., and Bovolenta, P. (2017). Adenohypophysis placodal precursors exhibit distinctive features within the rostral preplacodal ectoderm. Development 144, 3521–3532.

66. Schulte-Merker, S. (2002). Looking at embryos. In Zebrafish: A Practical Approach, (Oxford: Oxford University Press), pp. 39–40.

67. Sornson, M.W., Wu, W., Dasen, J.S., Flynn, S.E., Norman, D.J., O’Connell, S.M., Gukovsky, I., Carrière, C., Ryan, A.K., Miller, A.P., et al. (1996). Pituitary lineage determination by the Prophet of Pit-1 homeodomain factor defective in Ames dwarfism. Nature 384, 327–333.

68. De Souza, F.S.J., and Placzek, M. (2021). Conserved roles of Rax/rx3 genes in hypothalamus and pituitary development. Int. J. Dev. Biol. 65, 195–205.

69. Spoorendonk, K.M., Peterson-Maduro, J., Renn, J., Trowe, T., Kranenbarg, S., Winkler, C., and Schulte-Merker, S. (2008). Retinoic acid and Cyp26b1 are critical regulators of osteogenesis in the axial skeleton. Development 135, 3765–3774.

70. Takke, C., Dornseifer, P., v Weizsacker, E., and Campos-Ortega, J.A. (1999). her4, a zebrafish homologue of the Drosophila neurogenic gene E(spl), is a target of NOTCH signalling. Development 126, 1811–1821.

71. Toro, S., Wegner, J., Muller, M., Westerfield, M., and Varga, Z.M. (2009). Identification of differentially expressed genes in the zebrafish hypothalamus - pituitary axis. Gene Expr. Patterns 9, 200–208.

72. Wang, Y., Rovira, M., Yusuff, S., and Parsons, M.J. (2011). Genetic inducible fate mapping in larval zebrafish reveals origins of adult insulin-producing β-cells. Development 138, 609– 617.

73. Ward, R.D., Raetzman, L.T., Suh, H., Stone, B.M., Nasonkin, I.O., and Camper, S.A. (2005). Role of PROP1 in Pituitary Gland Growth. Mol. Endocrinol. 19, 698–710.

74. Wittkowski, W. (1986). Pituicytes. Astrocytes 1, 173–208.

75. Wittkowski, W. (1998). Tanycytes and pituicytes: morphological and functional aspects of neuroglial interaction. Microsc. Res. Tech. 41, 29–42.

76. Wu, W., Cogan, J.D., Pfäffle, R.W., Dasen, J.S., Frisch, H., O’Connell, S.M., Flynn, S.E., Brown, M.R., Mullis, P.E., Parks, J.S., et al. (1998). Mutations in PROP1 cause familial combined pituitary hormone deficiency. Nat. Genet. 18, 147–149.

77. Yoshida, S., Fujiwara, K., Nishihara, H., Kato, T., Yashiro, T., and Kato, Y. (2018). Retinoic acid signalling is a candidate regulator of the expression of pituitary-specific transcription factor Prop1 in the developing rodent pituitary. J. Neuroendocrinol. 30, e12570.

78. Zhang, S., Cui, Y., Ma, X., Yong, J., Yan, L., Tang, F., Wen, L., and Qiao, J. (2020). Single-cell transcriptomics identifies divergent developmental lineage trajectories during human pituitary development. Nat. Commun. 1–16.

79. Zhao, Y., Mailloux, C.M., Hermesz, E., Palkóvits, M., and Westphal, H. (2010). A role of the LIM-homeobox gene Lhx2 in the regulation of pituitary development. Dev. Biol. 337, 313– 323.

